# Post Adversity Changes in Nigro-Striatal Dopamine: A Mechanism for Anxiety Induced Exacerbated Innate Repetitive Behaviors

**DOI:** 10.1101/2025.06.21.660223

**Authors:** Lior Givon, Lee Amado, Shahaf Edut, Oded Klavir

## Abstract

Anxiety exacerbates symptoms in various psychiatric disorders. In conditions such as obsessive-compulsive disorder (OCD) or Tourette syndrome, anxiety intensifies stereotypic and repetitive behaviors. Rodent self-grooming, a structured, repetitive innate behavior, serves as an effective rodent platform for studying these behaviors in neuropsychiatric research. Anxiety is also linked to altered functioning of the dopamine (DA) system, particularly within the substantia-nigra pars compacta (SNc), the main DA source to the dorsal striatum through the nigro-striatal pathway. Striatal modulation by DA signal also plays a complex role in repetitive behaviors and OCD-like symptoms, suggesting this system as linking anxiety to the induced exacerbation of repetitive behavior. In the present study, we observed several long-term effects of anxiety inducing foot shock on grooming behavior. Recordings single unit neuronal activity in the SNc revealed distinct response patterns related to grooming behavior with changes in the magnitude and timing following the shock treatment. Notably, DA neurons of different nigro-striatal pathways demonstrated different changes in different response pattern type units. DA neurons projecting to the dorsolateral striatum (DLS) showed increase, while those targeting the dorsomedial striatum (DMS) exhibited decrease in transient activity -suggesting a shift in cortico-striatal circuitry of behavioral control. These neural changes were correlated with the observed behavioral alterations following adversity. Furthermore, targeted stimulation of DLS-projecting DA neurons rescued the anxiety-induced behavioral effects, highlighting the critical role of the nigro-striatal pathway to the DLS in mediating the interaction between anxiety and repetitive behaviors, thus offering future direction for mitigation of relevant psychiatric symptoms.

## Introduction

Anxiety is known to exacerbate symptoms in a broad spectrum of psychiatric disorders ^1,2^. Repetitive behaviors for example, observed in disorders such as obsessive-compulsive disorder (OCD) and Tourette syndrome are frequently exacerbated during periods of heightened anxiety ^3–5^. The rodent’s self-grooming behavior is an innate behavior, comprised of a structured sequence of repetitive actions. It is the most frequent of the innate behaviors observed in mice, and due to its frequency and similarity to other human behaviors, it is commonly used as a measure of such repetitive behavior in health and in models of neuropsychiatric disorders ^6–8^ . Grooming is an important adaptive behavior, playing an essential role in hygiene and temperature regulation, however, it can become maladaptive and result in skin lesions or fur loss ^9^. The parallel human excessive repeated behaviors like hand washing or trichotillomania, can lead to follicle damage, skin harm, and further negative psychological effects ^7^. In animal studies of neuropsychiatric disorders such as OCD, self-grooming is often noted to be disorganized and exacerbated due to anxiety ^10–13^. Some researchers suggested self-grooming microstructure alterations as a reliable marker for anxiety ^11^, and numerous studies have documented the relationship between anxiety and altered grooming behaviors, ^6,14,15^. Despite these insights, the underlying neural mechanisms responsible for the exacerbation of grooming behavior in response to anxiety remain mostly unknown.

Dopamine (DA) signaling demonstrates a complicated relationship with compulsive behaviors. Traditional theory posits that hyperactive DA signaling in cortico-striatal areas is a main contributor for OCD symptoms ^16,17^. In accordance, multiple studies shown that OCD-like behaviors are associated with excessive dopaminergic activity in cortico-striatal pathways ^18–21^. SAPAP3, which is a genetic mutant mouse expressing monoamine metabolism elevation in striatal and cortical areas ^22^ and increased striatal DA receptor density ^23^, exhibit both increased anxiety and compulsive ^23^.

Recent studies showed a more nuanced view of hyperactive DA in repetitive behaviors. For instance it has been shown that different DA projections from the Basal Ganglia can both promote and suppress repetitive grooming behavior ^21,24^. Pharmacological studies in humans also show ambiguity regarding the role of DA in OCD. Antipsychotics acting as DA antagonists can both exacerbate or improve symptoms ^25,26^. Similarly, drugs acting as DA re-uptake inhibitors also demonstrate this duality ^16,27–30^ leading to both exacerbation or improvement of OCD symptoms. Importantly, in those studies drugs are delivered systematically and lack system specificity. This is important as some work including our own shows that the effect DA exert on compulsive-like behaviors is circuit-specific and that different midbrain dopaminergic projections can either promote or suppress grooming behavior ^21,24^. Studies focusing specifically on DA of the substantia nigra pars compacta (SNc) showed that SNc DA can mitigate OCD-like compulsive grooming ^21^, moreover, stress was shown to modulate SNc activity ^31^ and even specifically attenuate SNc DA neurons ^32^. Midbrain DA, specifically of the SNc, regulates behavior by balancing competing pathways. Specifically, different DA neurons populations innervating different striatal areas are also considered respective to specific functional roles in the cortico-basal ganglia-thalamo-cortical (CBGTC) action control circuits ^33^. Moreover, those different populations, show unique properties in respect to features such gene expression and baseline firing rate ^34,35^, depending on their striatal target. The dorsomedial-striatum (DMS) is part of the associative loop, which processes action-outcome (A-O) and drives goal-directed behavior, and the dorsolateral-striatum (DLS) is part of the sensory motor loop, which mediates habits and stimulus-response (S-R) behaviors ^36,37^.

Therefore, in order to test whether the exacerbation of repetitive behavior under anxiety can be a result of DA activity and gain a deeper understanding of the neural and behavioral mechanisms through which anxiety modifies repetitive behaviors we recorded and analyzed the activity of differently projecting DA neurons of the SNc in mice, during induced and spontaneous self-grooming occurring before and after anxiety eliciting adverse event.

## Materials and Method

### Animals

Through this research, we used both DAT-CRE and C57BL/6 mice at the age of 12-16 weeks. Animals were housed together (up to 4 per cage) at 22 ± 2°C under 12-hour reversed light/dark cycles.

23 C57BL Mice (14 males) were included in the initial behavioral experiments.

7 C57BL Mice and 12 DAT-CRE Mice (1 Female) were included in all the electrophysiological experiments.

10 DAT-CRE mice (4 Males) were included in the optogenetic experiment.

Mice were allowed water and laboratory rodent chow ad libitum. The experiments were approved by the University of Haifa Ethics and Animal Care Committee, and adequate measures were taken to minimize pain and discomfort.

### Behavioral Paradigm

In this research, we employed a previously established ^24,38^ induced grooming paradigm (See Figure 1A & Figure 7A). The experiments took place in a 15cm x 15cm arena. All the experiments included a baseline period followed by three consecutive measuring days after the manipulation. Additionally, each measurement began with a habituation period, followed by a series of trials. Trials consisted of a water drop (75μL) applied using computer activated pump and were followed by inter-trial-interval of spanning randomly (3-4 Minutes).

**Figure 1.**
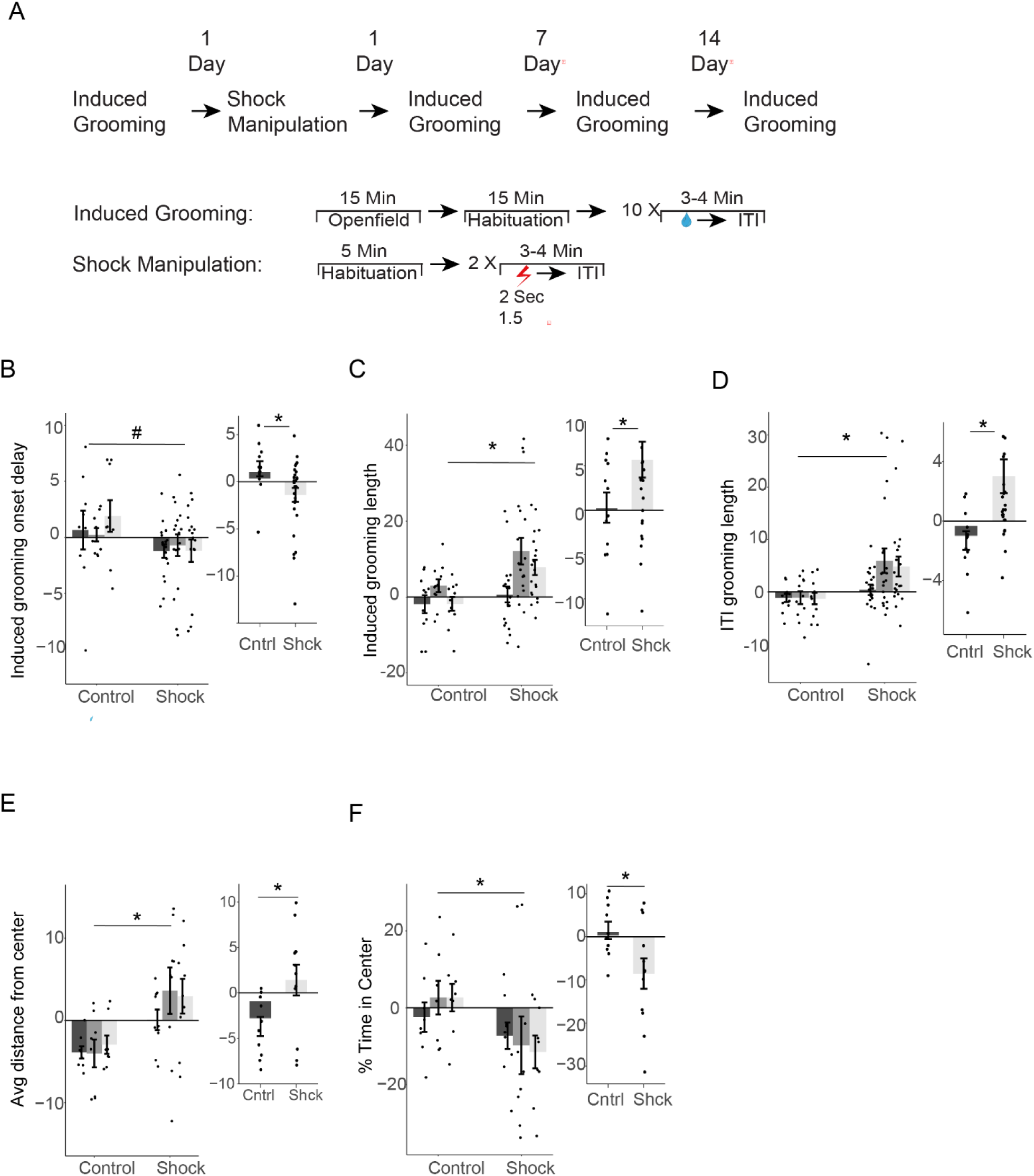
Changes in grooming behavior following shock exposure. **(A)** Illustration of the behavioral paradigms used to assess grooming behavior. The schematic also depicts the shock treatment administered to the shock group. **(B–D)** Grooming behavior scores (normalized by individual animal baseline) comparing control and shock groups. Large plots separate by measuring days following the shock-box; smaller insets display collapsed – by group values. **(B)** Delay in initiation of induced grooming bouts. **(C)** Duration of induced grooming bouts. **(D)** Duration of ITI grooming bouts. **(E–F)** Open field behavioral scores (normalized by individual animal baseline) comparing control and shock groups. Large plots separate by measuring days following the shock-box; smaller insets display collapsed – by group values. **(E)** Distance from the center of the open field. **(F)** Percentage of time spent in the central quarter of the open field. In each plot, bars represent means of corresponding measures with error bars showing ±SEM. Asterisks indicate significant post-hoc comparisons (*, p<.05; **, p<.01; ***, p<.001). Pound signs indicate non-significant trends (#, .05<p<.1).

### Shock Manipulation

Twenty-four hours after the baseline measure the animals went through an electrical shock shown to induce anxiety-like behavior in the openfield test. To induce a non-associative anxiety, we used a previously established procedure ^38,39^. Following 5 minutes of habituation time, animals were exposed to two strong electrical foot shocks (2s, 1.5 mA). The delay between the shocks was 40s. Shocks were applied in a Med-Associates operant conditioning chamber (Med-Associate, USA). Control animals were placed in the chambers for the same time duration, without the electrical shocks.

### Open-field

To measure anxiety, mice were placed in in an open field (50cm x 50cm) before each measurement, for a total of 15 minutes. The open field is widely used as a method to evaluate anxiety in rodents. Anxious rodents usually exhibit thigmotaxis, spending more time near the walls and less time in the center, hence open field centrality is used as a common marker for anxiety ^40^. To overcome possible confounds, centrality was assed according to two measures:

1. Average Distance – we calculated the center point and measured the distance of the animal to the center point in each frame and calculated the mean distance.
2. Center Quarter – We counted the number of frames in which the animal was closer to the center point than to any of the corners, thus defining a ’Center Quarter’.

Surgical procedures:

Surgery: Animals were sedated using IP injection of ketamine (80mg/kg) and xylazine (8mg/kg) and anesthetized with 1-2% isoflurane modified during procedure according to reflexes checks. After full anesthesia, the head was fixed in a stereotactic device and an injection of the analgesic

NoroCarp (20mg/kg) was delivered. The body temperature kept at 37 Celsius via a heating pad. Skin was disinfected using ethanol and betadine and washed with saline before incision. For three days following the procedure, animals were kept under close supervision and received daily treatments of NoroCarp (0.01 ml/20g) and Amoxicillin (2.5 ml/kg).

Injections: To test the role of DA projections to the DLS we used two injections of pAAV-EF1a-double-floxed-hChR2(H134R)-EYFP-WPRE-HGHpA (AAV Retrograde) in the same hemisphere at two depths for maximal cover: (AP: 0.5, ML: +/-0.5; DV: −2.5, −2.7; total 0.45μL in each injection). The expression of the ChR2 opsin allowed us to identify DA neurons projecting to the DLS using phototagging. pAAV-EF1a-double floxed-hChR2(H134R)-EYFP-WPRE-HGHpA was a gift from Karl Deisseroth (Addgene plasmid # 20298 ; http://n2t.net/addgene:20298 ; RRID:Addgene_20298). To target DA projections to the DMS we used the same vector, again with two depths injection for maximal cover (AP: 0.5, ML: +/-2.5; DV: −2.8, −3.0.; 0.45μL each injection).

To optogenetically activate DA projections to the DLS we used two injections of pAAV-hSyn-DIO-somBIPOLES-mCerulean (AAV-Retrograde) in the both hemispheres at two depths for maximal cover (AP: 0.5, ML: +/-2.5; DV: −2.8, −3.0.; 0.45μL each injection). hSyn-DIO-somBiPOLES-mCerulean was a gift from Simon Wiegert (Addgene plasmid # 154951 ; http://n2t.net/addgene:154951 ; RRID:Addgene_154951)^42^.

Cannula implant: To drop water and we implanted 0.819mm diameter cannula above the mice’s forehead.

Optrodes: To record neaural activity, we used a custom-made optrodes, implanted three weeks after the injection surgery. The optrode was built using a Milmax connector (Mill-Max, USA) sliced to a length of 18 pins. Tungsten wires (Wiretronics, Sweden) were threaded through 8 pins from the left side and 8 pins from the right side, with a ground cable threaded through the middle. An optical fiber (Thor-Labs Inc, USA) was glued to the optrode using epoxy glue to allow for light stimulation above the recording site. After grounding the optrode to the skull by connecting the ground cable with a tiny screw, it was positioned in one hemisphere of the SNc (AP: −3.15, ML: +/-1.25, DV: 4.5) and secured using C&B Metabond (Parkell, USA) quick-drying glue, and dental cement

Optical Fibers: To optogentically activate the DLS projections to the SNc we implanted bilateral optical fibers (Thor-Labs Inc, USA). Fibers were positioned in both hemispheres of the SNc (AP: −3.15, ML: +/-1.5, DV: 4.1).

### Electrophysiological recording and light stimulation

performed using Neuralynx software (Neuralynx, USA). The signal was filtered (600-6,000 Hz) and amplified with a headstage. Recorded data was sorted manually by Plexon’s Offline Sorter (Plexon, USA), and analyzed with Python 3.0, utilizing NeuralynxIO library.

Light intensity: The light power delivered to the target tissue was calibrated to 15 mW at the fiber tip, accounting for light loss in the optical fiber, which was sufficient for effective opsin activation.

Light stimulation in photo-tagging: To determine the light response of recorded units, we used a photo-tagging protocol consisting of two 150 single light stimulations, 0.5 Hz trains, (3ms on and 1997 ms off) for 5-minute duration each. We utilized a moving window approach to analyze the light-responsiveness. We collected all light events and utilized a series of t-tests to compare the average baseline neural activity [-30ms, −1ms] to successive time windows after the light stimulation. The size of the moving window was set to 5ms, with the initial window spanning [0ms, 5ms]. We searched for a time-window with significantly higher activity compared to the baseline. If a given window did not yield a significant difference from baseline, we advanced the window onset by 3ms and repeated the analysis, keeping the window size at 5ms. This process continued until reaching the 15ms. Hence, we tested all the following time windows: [0-5ms], [3-8ms], [6-11ms], and [9-14ms]. This approach takes into account a possible late response to the light stimulation. We validated the results of this statistical approach by comparing PSTHs of light-responsive vs non-light responsive units.

Light stimulation in behavioral experiment : Light stimulation was employed using the BiPOLS tool^42^ which incorporates two distinct opsins. For the inhibitory condition, a continuous inhibitory light signal was delivered for a duration of 2 seconds. The excitatory condition involved a 25 Hz stimulation frequency applied for 2 seconds.

### Histology

All animals had undergone perfusion. Brain slices and staining, and brain tissues were examined under a light microscope to confirm the presence of the appropriate fluorophore, verifying successful viral transduction in the targeted regions.

Perfusion: Animals were anesthetized using an IP injection of pentobarbital (0.04ml/20g). After being fully anesthetized (verified by reflex check) cold phosphate buffered saline (PBS, pH 7.4) passed through the heart, followed by 4% solution of PFA (Paraformaldehyde). After decapitation, heads were placed in a PFA solution for 72 hours. Extracted brains placed in the same solution for additional 24 hours and then placed in a 20% sucrose solution. All solutions were stored in a 4°C.

Brain Slicing and Staining: Using Leica’s microtome (Leica Biosystems, Germany) coronal sections sliced (40μm) and mounted. Sections were either immediately mounted on gelatin-coated slides or stored in cryoprotectant solution (25% glycerol, 30% ethylene glycol in PBS, pH 6.7) at −20°C until further use. Prior to covering, DAPI (4’,6-diamidino-2-phenylindole; Sigma-Aldrich, USA) was applied directly to the glass slides containing the mounted sections.

Microscopy: Axio Imager 2.0 fluorescent microscope (Carl Zeiss AG, Germany) was used to validate and photograph the brain slices.

### Data Analysis

All measures regarding the animals’ behavior were detected and extracted using DeepLabCut ^42^ and custom-made algorithms were used to extract grooming from spatial location and analyze it accordingly. In each trial we measured the percentage of time that the animal spent grooming, the delay in induced grooming initiation after the drop and the average length of a grooming bout.

Behavioral data handling: We observed considerable variability in grooming behavior across animals, including baseline differences and individual grooming tendencies. To account for this and enable standardized comparisons, each behavioral measurement was normalized to the corresponding baseline value from the matching trial (e.g, 1_𝑇𝑟𝑖𝑎𝑙1𝐺𝑟𝑜𝑜𝑚𝑖𝑛𝑔%_ = 1_𝑇𝑟𝑖𝑎𝑙1𝑔𝑟𝑜𝑜𝑚𝑖𝑛𝑔%_ − 𝐵𝑎𝑠𝑒𝑙𝑖𝑛𝑒_𝑇𝑟𝑖𝑎𝑙1𝐺𝑟𝑜𝑜𝑚𝑖𝑛𝑔%_), following the approach of a previous study ^24^. This normalization allowed us to assess post-manipulation behavior relative to each animal’s baseline. For each measurement day, we averaged all trials, then computed the mean across all days. Final scores for each dependent variable were calculated by averaging the normalized trial data per measurement, followed by averaging across the three measurements to quantify the overall effect of the manipulation per animal:

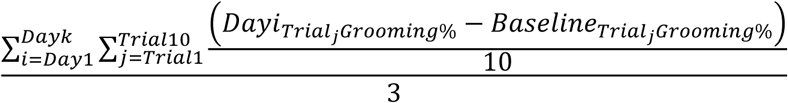

Physiological data handling:

Metrices for comparing firing patterns (Figure2):

1-Magnitude: for transient peak response − The magnitude corresponds to ratio between the maximal FR bin (250ms) observed within the peak window ([-1000,1000]), divided by the designated baseline for each event. This metric can be expressed by the following formula:

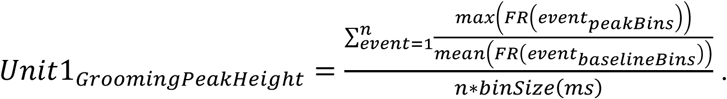

Magnitude: for stable response, corresponds to the mean FR after the event divided by the mean FR before the event.

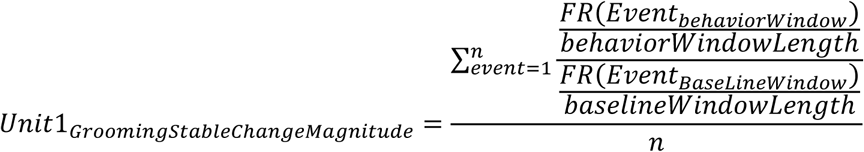

2- Onset timing refers to the temporal position of the maximal FR bin within the peak window can be expressed as follows:

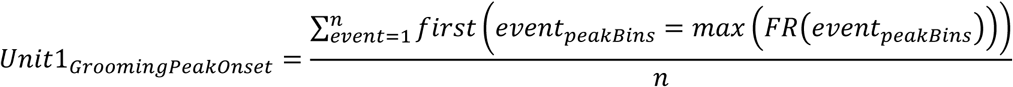

### Statistical Analysis

We conducted ANOVAs using the aov() function from the **stats** package, or Welch ANOVAs via aov_ez() from the **ez** package when variances were unequal. T-tests (t.test() from **stats**) were used both for comparisons between two groups and for pairwise comparisons among multiple groups when applicable. Dunnett’s post-hoc tests were conducted using glht() from the **multcomp** package. Trials with missing data (due to animal deaths, recording issues, or pump blockages) were excluded, and outliers (defined as values exceeding ±3 standard deviations from the mean) were removed; when analyzing multiple groups, outliers were identified and removed within each group separately. Welch’s t-test or Welch ANOVA was applied when variances were unequal, and sphericity was tested prior to ANOVA. To assess correlations, we used linear regression (lm()) and Chi-square tests for proportions were conducted using the prop.test() function from the stats package. Graphs were plotted using **ggplot** package. Statistical significance was defined as *p* < 0.05.

## Results

### Significant, unexpected and inescapable foot shock induces long term changes in grooming behavior as well as anxiety related behavior

To test the long-term behavioral effects of the shock manipulation we compared grooming behavior (externally and spontaneously induced) before and at three time points after the shock manipulation (Figure 1A). We specifically considered three behavioral markers shown to involve the DA system and to be modulated by a fear eliciting cue (Givon et al., 2024). First, initiation delay - the time required for the onset of grooming behavior in response to the external grooming induction cue, was shortened by the adverse experience and remained short, long after the shock (Figure 1B). Indicating maintenance of a faster reaction time in the shock group (M= −1.04, SD=3.54) compared to the control group (M = 0.93, SD=3.9), long after the manipulation. Mixed ANOVA model with Shock as a between-subjects factor and Measurement as a repeated within-subjects factor revealed no significant main effect of Measurement (F(2,48) = 0.14, p = 0.868) and no Shock × Measurement interaction (F(2,48) = 0.74, p = 0.483), indicating that delay patterns remained stable across time and did not differ by group across measurement epochs. A trend-level main effect of shock was observed (F(1,24) = 2.97, p = 0.098), suggesting potential differences in grooming initiation latency between the shock group and the control group. Given the absence of interaction, we again assessed the main effect of Shock averaged across all measurements. This analysis revealed a significant difference in initiation delay between groups, with animals in the shock group exhibiting significantly shorter delays relative to controls (t(38) = 2.24, p < 0.05). These findings suggest that shock exposure reduces the reaction time to grooming inducing cue.

We next considered the long-term effects of the shock manipulation on the length of the induced grooming bout and found that it became longer (Figure 1C) in the shock group (M=6.83 SD=12.19) compared to the control group (M=0.24 SD=6.74). A Type III Mixed ANOVA with welch-correction found a main effect for group indicating that the increase in the shock group was significant (F(1,28)=6.34, p <0.02). A significant main effect of Measurement was observed (F(1.81,50.77) = 6.98 p < 0.003), indicating grooming bout length varies with time. However, there was no significant Group × Measurement interaction (F(1.81,50.77) = 1.62, p = 0.209), suggesting that group differences did not vary across time. Given the lack of interaction, we again examined the main effect of shock, averaged across all timepoints. Welch’s corrected t-test revealed a significant difference in the length of the induced bout (t(35.28) = –2.04, p < 0.05), with animals in the experimental group exhibiting longer duration of grooming bouts. These findings indicate that the adverse experience led to a sustained increase in the duration of induced grooming bouts.

Not just the externally induced grooming bouts were prolonged due to the shock experience but also the length of spontaneous grooming bouts, initiated during the ITI (Figure 1D). Welch corrected Mixed ANOVA revealed a significant main effect of shock (F(1,29) = 5.71, p < 0.025), with longer ITI grooming observed in the shock group (M=3.71, SD=8.43) compared to the control group (M=3.38, SD=3.38). There was no significant main effect of Measurement (F(1.77,51.31) = 2.25, p = 0.122), or of Shock × Measurement interaction (F(1.77,51.31) = 2.27, p = 0.119). Given the lack of interaction, we again examined the main effect of shock, averaged across all time points. Welch’s t-test revealed a robust group difference (t(37.49) = –3.32, p < 0.002), as animals in the shock group exhibited significantly longer ITI grooming duration relative to controls.

To test whether the long-term effects of the shock manipulation on measures of general anxiety, before each measuring day we tested the mice in an open field and found that indeed mice still showed measures of anxiety as far as three weeks post adversity. Animals in the shock group (M=44.29, SD=128.19 maintained a significantly greater average distance from the center (Figure 1E) relative to controls (M=-72.47, SD=68.05). Welch corrected Mixed ANOVA revealed a significant main effect of shock (F(1, 15) = 6.46, p < 0.025) with no main effect of Measurement (F(1.54, 23.06) = 2.14, p =0.149) and no Shock × Measurement interaction (F(1.54, 23.06) = 1.64, p =0.217), indicating an increase in anxiety dependent thigmotactic behavior. The lack of significant interaction suggests that the behavioral pattern is stable across the testing session. Therefore, the average effect of shock exposure across all the measures was evaluated. This analysis also revealed a significant main effect of shock (t(20) = 2.44, p<0.025).

The effects of anxiety in the open field were also evident from the time the mouse spent in the central quarter of the arena which was also decreased in the shock group (Figure 1F) compared to the control group (M=-0.1, SD =0.11 vs M=0.01, SD=0.16). A Mixed ANOVA shown a marginally significant effect of shock (F(1,15) = 4.27, p = 0.056) with no main effect of Measurement (F(2,30) = 0.05, p = 0.96) and Measurement × Shock interaction (F(2, 30) = 0.80, p = 0.461). Similar to the center distance. A complementary analysis averaging all the shock measurements again revealed a significant main effect of shock (t(20) = 2.35, p < 0.03), reinforcing the impact of shock on center-avoidant behavior. Together these two open field measures suggest that the shock manipulation induces long term anxiety.

### SNc neurons produce several types of grooming firing patterns, differently responding to adversity

To characterize SNc neural activity during grooming we recorded extracellularly in the SNc of mice during the behavioral paradigm. The spiking activity of each recorded unit around both the initiation and the termination of each grooming bout was examined and responding units were statistically identified according to the type of their response. The average firing rate during a set baseline period (five seconds to two seconds prior to the event) was compared to the average firing rate immediately following the behavioral event (the four seconds following the event), for grooming initiation or termination separately, using a paired t-test across all grooming events. Units reaching statistical significance (p < 0.05) were determined as responsive. For significant units, if the mean firing rate exceeded the baseline after the event, units were classified as “Stable Excitation” units, indicating excitatory response to the onset. Units who decreased below baseline on the other hand, were classified as “Stable Inhibition” units, indicating inhibitory response to the onset. Those units with sustained response both to grooming initiation and to grooming termination constituted a bit over quarter of the recorded neurons (Figure2A). Another type of response was also identified which was brief and occurred immediately with the behavioral event was termed Transient Peak response. To find this response we used the same baseline window, but the response window was divided into a peak window (the two seconds around the event) and a post-peak period (the two seconds following the peak window). Firing rate within the peak window had to be significantly higher than the baseline period but also higher than the post peak period to be considered a Transient Peak response. Our categorization was mutually exclusive as each unit is assigned to either a single response type or none therefore in cases where a unit had a stable response following the peak, it was still counted as Transient Peak (Figure2A).

The categorization was a mutually exclusive scheme in which each unit is assigned to either a single response type or none. If a unit had a stable response in addition to a peak, it was still counted as Transient peak, as the stable response can be counted as a following sustained response. (Figure2A).

We next characterized the response patterns of the different response types. To get a full picture of full grooming bout response types, we tested the distribution of the different grooming termination types within each grooming initiation type. Next, we examined whether there is a difference in the response of each neuronal type before and after the shock manipulation, which could be related to the behavioral effects observed following the shock. To compare the neural responses, we used two metrics: the magnitude of the neural response and the timing of the response onset relative to the grooming event (see Methods section for formulas).

Grooming initiation stable excitation units, almost exclusively responded to termination in stable inhibition (Figure 2B left ; χ²(3) = 149.79, p < 0.001; residual = 11.76), creating the first type of grooming dependent neuronal response - Stable excitation firing pattern (Response scheme on Figure 2B right). This type of response (Example firing pattern see Figure 2C) showed increased magnitude and delayed response after the shock manipulation (Figure 2D). Looking at grooming onset Welch’s *t*-test revealed a significant increase in magnitude following the shock, with pre-shock units showing lower excitation (M = 1.49, SD = 0.41) compared to post-shock units (M = 3.60, SD = 4.36), *t*(213.27) = −6.77, *p* < 0.0001). Comparing the timing of excitation onset before and after the shock with an independent samples t-test indicated a significantly delayed onset post-shock (M = 136.10, SD = 131.00) relative to pre-shock (M = 91.64, SD = 124.00), *t*(255) = −2.26, *p* < 0.025).

**Figure 2.**
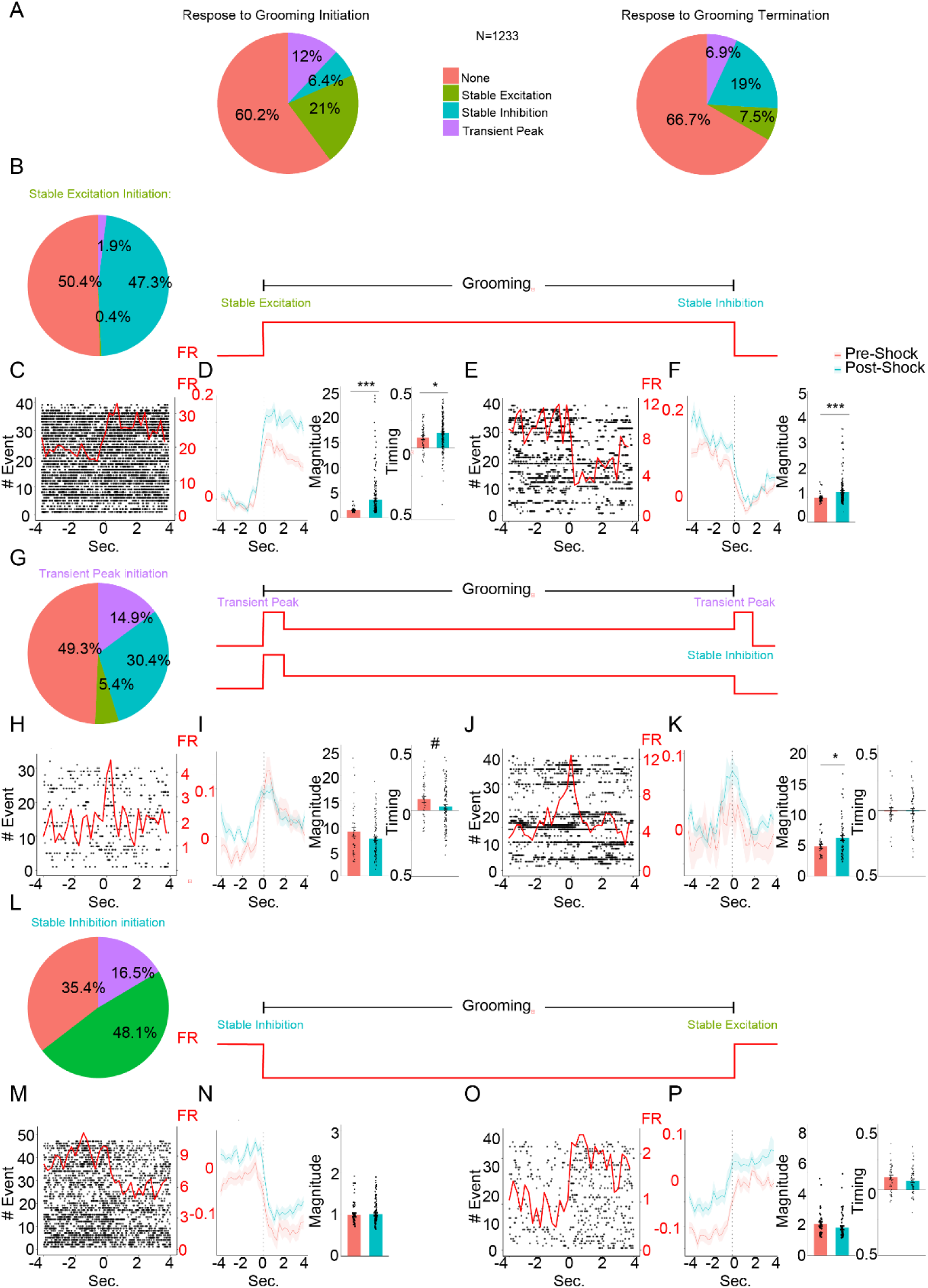
Neural response types to grooming initiation and termination, and their modulation following shock exposure. **(A)** Pie charts showing the distribution of the three neural response types to grooming initiation (left) and grooming termination (right). **(B) Left**: Distribution of grooming termination response types among units that exhibited stable excitation at grooming initiation. **Right**: Response scheme of grooming dependent Stable excitation firing pattern **(C)** Example firing pattern of Stable excitation type around grooming onset **(D) Left:** Peristimulus time histogram (PSTH) of all stable excitation units responding to grooming initiation, before and after shock. **Middle**: Bar plot showing a significant increase in response magnitude. **Right**: Bar plot showing a shift toward earlier response timing after shock. **(E)** Example firing pattern of Stable excitation type around grooming termination **(F) Left:** Peristimulus time histogram (PSTH) of all stable excitation units responding to grooming termination **Middle**: Bar plot showing a significant increase in response magnitude. **Right**: Bar plot showing no difference in response timing after shock. **(G) Left**: Distribution of grooming termination response-types among transient peak units at grooming initiation. **Right**: Response scheme of grooming dependent transient peak firing patterns **(H)** Example firing pattern of transient peak type around grooming onset **(I) Left:** Peristimulus time histogram (PSTH) of all transient peak units responding to grooming initiation, before and after shock. **Middle**: Bar plot showing no significant change in response magnitude. **Right**: Bar plot showing a trend toward earlier response timing following shock. **(J)** Example firing pattern of transient peak type around grooming termination **(K) Left:** Peristimulus time histogram (PSTH) of all transient peak units responding to grooming termination **Middle**: Bar plot showing a significant increase in response magnitude. **Right**: Bar plot showing no difference in response timing after shock. **(L) Left**: Distribution of grooming termination response types among units that exhibited stable inhibition at grooming initiation. **Right**: Response scheme of grooming dependent Stable inhibition firing pattern **(M)** Example firing pattern of Stable inhibition type around grooming onset **(N) Left:** Peristimulus time histogram (PSTH) of all stable inhibition units responding to grooming initiation, before and after shock. **Middle**: Bar plot showing no significant change in response magnitude. **Right**: Bar plot showing no change in response timing after shock. **(O)** Example firing pattern of Stable inhibition type around grooming termination **(P) Left:** Peristimulus time histogram (PSTH) of all stable inhibition units responding to grooming termination **Middle**: Bar plot showing no significant change in response magnitude. **Right**: Bar plot showing no difference in response timing after shock. In each plot, bars represent means of corresponding measures with error bars showing ±SEM. In peri-event histograms, each line is representing the overall average of the corresponding measure with shaded ±SEM. Asterisks indicate significant comparisons (*, p<.05; **, p<.01; ***, p<.001). Pound signs indicate non-significant trends (#, .05<p<.1).

Termination stable inhibition units (Example firing pattern see Figure 2E) the termination response of initiation stable excitation also showed decreased magnitude of inhibition (higher firing rate) after the shock manipulation though the timing of the termination response did not change. (Figure 2F). Looking at grooming offset Welch’s *t*-test revealed a decreased inhibition magnitude in the post-shock measurements (M=1.22, SD=0.58) compared to pre-shock (M=0.97,SD=0.17), (*t*(222.5) = −4.92, *p* < 0.001). Comparing the timing of inhibition onset before and after the shock with Welch t-test indicated no significant difference between the pre-shock (M = −333.91, SD = 191) and post-shock (M = −356.47, SD = 148), (t(83.71) = 0.82, p =0.412).

Grooming initiation transient peak units, responded to termination in either stable inhibition or transient peak (Figure 2G left ; χ²(3) = 31.34, p < 0.001; Stable Inhibition residual = 3.55; transient peak residual = 3.83), creating the second type of grooming dependent neuronal response – transient peak firing pattern (Response schemes on Figure 2G right). This type of response (Example firing pattern see Figure 2H) showed a trend for an earlier response onset following the shock but no change in the magnitude around grooming onset (Figure 2I).

A Welch’s *t*-test showed no significant difference in spike height between pre-shock (M = 8.81, SD = 5.37) and post-shock conditions (M = 7.44, SD = 3.63), *t*(46.33) = 1.43, *p* = 0.159). A t*-*test comparing spike onset timing between pre-shock (M = 94.7, SD = 161.18) and post-shock measurements (M = 32.4, SD = 175.73) revealed a marginally significant difference, *t*(140) = 1.88, *p* = 0.063). Unlike the initiation transient peak, the termination transient peak units (Example firing pattern see Figure 2J), did show an increased magnitude after the shock manipulation with no change in the timing of the termination response. (Figure 2K). Looking at grooming offset t-test revealed a higher peak height following the shock (M=7.67, SD=3.94) compared to the pre-shock (M=5.98, SD=2.18) with Welch t-test (*t*(75.88) = −2.49, *p* < 0.015). However, t-test revealed no change in timing (*t*(81) = −0.02, *p* = 0.983).

Both the grooming bout length and the magnitude of grooming termination transient peak were found to be affected by the shock manipulation. Looking at the relations between the two parameters, we found that the height of the transient peak predicted the magnitude of the behavioral effect. A weighted linear regression was conducted to examine the relationship between the magnitude of the termination responses and the normalized length of the induced grooming bout (Supp. Figure 1A), with the number of responding transient peak units used as weights. The model revealed a significant positive association (r=0.53, p < 0.037, *R*² = 0.28, adjusted *R*² = 0.22), similar analysis showed even stronger relations between the termination response magnitude and the normalized average length of spontaneous bouts measured during the ITI (*r=0.65*, *p* < 0.01, *R*² = 0.42, adjusted *R*² = 0.39). This suggests that the magnitude of the termination response of transient peak units is a consistent predictor of the length of grooming behavior, possibly due to increased threshold required for termination of grooming following the shock. The height of termination response magnitude expectedly did not correlate with the normalized delay (*β* = 0.13, *SE* = 0.28), *t*(14) = 0.44, *p* = 0.666, *R*² = 0.014), indicating that termination spike amplitude does not predict latency in grooming onset. As there was also a significant increase in inhibition magnitude following the shock, we conducted the same correlation with this type of grooming termination response pattern which yielded no significant correlation with either induced grooming bout length (*r=0.0265*, *p* = 0.193) or spontaneous grooming bout length (r=0.332, p = 0.090).

Grooming initiation stable inhibition units, almost exclusively responded to termination in stable excitation (Figure 2L left ; χ²(3) = 211.89, p < .001; residual = 13.75), creating the third type of grooming dependent neuronal response - Stable inhibition firing pattern (Response scheme on Figure 2L right). This type of response (Example firing pattern see Figure 2M) showed no difference in magnitude or timing of the response after the shock manipulation (Figure 2N). Looking at grooming onset *t*-test revealed non-significant in magnitude following the shock, comparing pre-shock units (M = 0.99, SD = 0.22) and post-shock measurements (M = 1.01, SD = 0.26), *t*(74) = −0.34, *p* = 0.74). Comparing the timing of inhibition onset before and after the shock with an independent samples t-test indicated no significant difference between the pre-shock (M = −373.68, SD = 227) and post-shock (M = −398.19, SD = 258), (t(74) = 0.40, p = 0.691).

Termination stable excitation units (Example firing pattern see Figure 2O) also showed no change in increased magnitude or timing after the shock manipulation. Looking at grooming offset t-test revealed no significant difference in magnitude between groups, *t*(87) = 1.32, *p* = 0.190) or in timing of the excitation (*t*(87) = 0.70, *p* = 0.488).

### Differences in electrophysiological properties of DLS vs DMS projecting DA neurons

To test for unique characteristics of differentially projecting DA neurons we injected retrograde AAV carrying Cre-dependent ChR2 into either the DLS or the DMS of DAT-Cre mice. Opsin expressing DA neurons (For example see Figure 3A Top) were recorded by implanting an optrode in the SNc (For electrode placement scheme see Figure 3A bottom) and photo-tagged to identify their projection target (For projection scheme see Figure 3B - for full details see the Methods). Given the specific targeting of retrograde-dependent expression in DA which depends on the exact location and volume of injected tissue, the number of light-responsive units was limited. Of the units recorded in animals injected with the retrograde viral vector into the DLS, 47 units (13.9%) were photo-tagged (expression and photo-tagging examples Figure 3C Top; distribution on Figure 3C Bottom) while 33 units (5.7%) were photo-tagged among those recorded from animals injected with the retrograde viral vector into the DMS (expression and photo-tagging examples Figure 3D Top; distribution on Figure 3D Bottom). As DA neurons are characterized by relatively unique properties such as their firing rates (2-10 HZ) and the relatively stable and reliable course of action potential ^43^, these properties were used to assess the differences between the photo-tagged units and the non-phototagged units in each projection tagging experiment (DLS or DMS).

**Figure 3.**
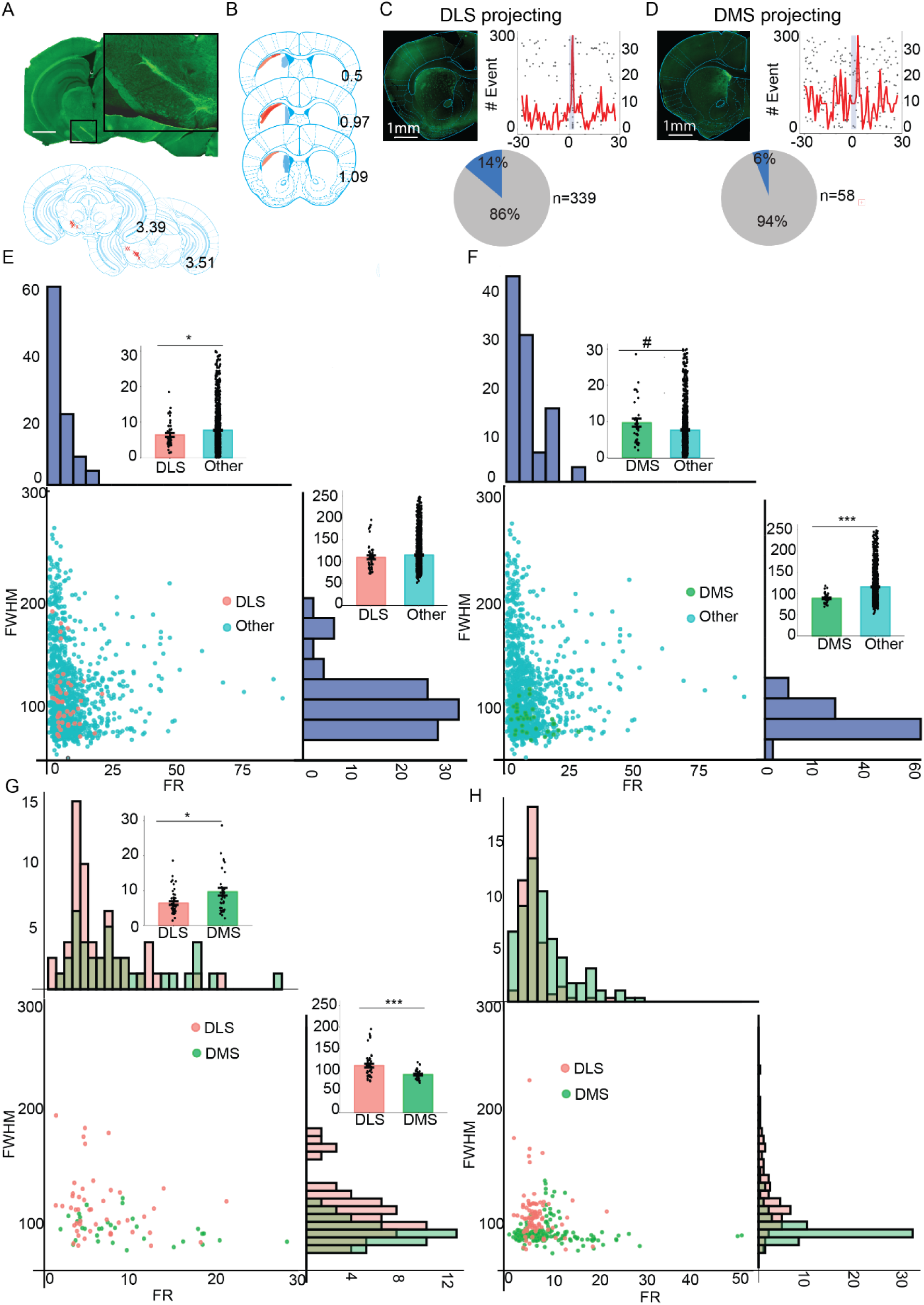
Identification and comparison of dopaminergic SNc units projecting to the DLS and DMS. **(A)** Top: representative coronal section showing the expression in SNc DA neurons. Bottom: anatomical target schemes showing the SNc. X’s mark histologically confirmed optrode positions from all animals included in the experiment. **(B)** An overlaid, histologically confirmed viral expression areas at the DLS (in shades of red) and DMS (in shades of blue) from all included animals. **(C)** Top left: Representative coronal section showing the expression of SNc DA neuronal projections in the DLS. Top right: A raster plot overlaid with PSTH of photo-tagged SNc DA unit projecting to the DLS. Bottom: Percentage of light-responsive DLS-projecting units recorded in the experiment. (Top left: Representative coronal section showing the expression of SNc DA neuronal projections in the D Top right: A raster plot overlaid with PSTH of photo-tagged SNc DA unit projecting to the D Bottom: Percentage of light-responsive D S-projecting units recorded in the experiment.**( )** Comparison of DLS-phototagged units to non-DLS units. Scatter plot shows individual units by firing rate and waveform width (FWHM). **Top and right**: Histograms display distributions of firing rate and FWHM. **Next to histograms**: Bar plots show a significantly higher firing rate in DLS units compared to non-DLS units, with no significant difference in FWHM. (Comparison of DMS-phototagged units to non-DMS units. Scatter plot shows individual units by firing rate and FWHM. **Top and right**: Histograms display distributions of firing rate and FWHM. **Next to histograms**: Bar plots show a trend toward higher firing rate in DMS units compared to non-DMS units, and a significant shorter FWHM in the DMS units. **( )** Direct comparison of DLS- and DMS-phototagged units. Barplots show higher firing rates in DMS units and broader waveforms (higher FWHM) in DLS units. (Scatter plot and histograms summarizing firing rate and waveform differences comparing emulated DLS- and DMS-photoagged units. In each plot, bars represent means of corresponding measures with error bars showing ±SEM. In peri-event histograms, each line is representing the overall average of the corresponding measure with shaded ±SEM. Asterisks indicate significant comparisons (*, p<.05; **, p<.01; ***, p<.001). Pound signs indicate non-significant trends (#, .05<p<.1).

We first assessed the differences in firing rate between DLS-projecting DA neurons and the general population (Figure 3E). We found that firing rates in photo tagged in DLS-projecting DA units were significantly lower than the general population of recorded units, but no significant difference was found in the width of the recorded action potential. Welch t-test revealed a significant difference in firing rates in DLS-projecting units (M = 6.40, SD = 3.64) compared to the general population (M = 7.74, SD = 6.63), (t(57.60) = −2.34, p < 0.02). Full Width at Half Maximum (FWHM), which reflects the duration of the signal at 50% of its peak amplitude was however non-significant when comparing the DLS-projecting neurons (M = 110, SD = 29.7) and the general population (M = 115, SD = 39.2) using Welch t-test (t(52.60) = −1.16, p = 0.2505). We next assessed the differences in firing rate between DMS-projecting DA neurons and the general population (Figure 3F). We found that firing rates in photo tagged in DMS-projecting DA units tended to be higher than the general population of recorded units, and that the width of the recorded action potential (spike) in those tagged neurons was significantly shorter than the general population. T-test revealed a trend toward a higher firing rate in DMS-projecting neurons (M = 9.66, SD = 6.39) compared to the general population (M = 7.64, SD = 6.55), (t(1203) = 1.75, p = 0.0803) and a Welch t-test revealed a significant difference in FWHM, with DMS-projecting neurons (M = 88.8, SD = 12.2) showing lower durations than the general population (M = 116.0, SD = 39.1), (t(52.71) = −11.18, p < 0.0001).

While the SNc is considered one of the main nuclei of DA neurons in the brain ^44^ with a majority of DA neurons ^45^. The DMS and the DLS photo-tagged DA units entailed electrophysiological differences from the general SNc neural population which includes also a large GABAergic and Glutamatergic subpopulations ^45^. As DA neurons are known to differ in various properties based on the brain region they project to ^34,35^, we next continued to comparing both populations directly. Past studies comparing DLS and DMS-projecting DA neurons demonstrated biophysical properties and wiring differences ^46^. Examining the differences in the firing properties between the two neurons populations directly indeed demonstrated that DLS-projecting DA neurons produce significantly lower firing rates and a prolonged action potential course compared to DMS-projecting DA neurons (See Figure 3G). Welch t-test showed significantly lower firing rates of DLS-projecting neurons (M = 6.40, SD = 3.64) than DMS-projecting neurons (M = 9.66, SD = 6.39), t(46.80) = −2.64, p < 0.015) and a significantly longer FWHM of DLS-projecting neurons (M = 110.0, SD = 29.7) than DMS-projecting neurons (M = 88.8, SD = 12.2), t(65.32) = 4.38, p < 0.001).

Based on the observed differences between the two photo-tagged populations, we classified the all-recorded units from all the animals (See Figure 3H). Units were designated as DLS-projecting (n=114) if they were either photo-tagged or fell within −0.5 and + 0.25 standard deviations of the mean FR of the photo-tagged DLS units and within ±0.5 SD of the mean FWHM. Similarly, units were classified as DMS-projecting (n=173) if they were either photo-tagged or fell within −0.25 SD and +0.5 SD of the mean FR of the DMS and within ±0.5 SD of the mean FWHM. We used 0.25 SD on the two ends of the firing rate ranges to avoid overlapping between the two groups of classified units. These differences were used to emulate projection separation on all units using photo-tagging classification data.

### Grooming firing patterns types of DA neurons are a function of their projection target and the exposure to adversity

Assessing the proportional firing pattern types responding to grooming (Transient Peak, Stable Excitation or Stable Inhibition) in DLS and DMS projecting photo-tagged DA units and in the non-tagged neural populations revealed that while the distribution of different pattern types in DMS projecting DA units are no-different than the general population, DLS projecting units are uniquely distributed between the different types, and seem to have a significantly larger portion of transient peak firing pattern (Figure 4A - top). A chi-squared test showed a significant difference in transient peak neurons proportions across the three groups( χ²(2) = 19.20, *p* < 0.001) with post hoc comparisons comparing the DLS and DMS projecting photo-tagged neurons to the non-tagged control group with Holm adjustments, revealing a significant difference between the non-tagged control group and the DLS-tagged group (χ² = 15.98, *p* < 0.001; adjusted *p* = 0.001), but no significant difference between the control and DMS groups (χ² = 0.45, *p* < 0.50 ;adjusted *p* = 0.50). A similar proportional analysis for the stable excitation or the stable inhibition pattern type neurons did not reveal any significant difference (χ²(2) = 2.86, *p* =0.24 and χ²(2) = 0.39, *p* =0.82, respectively).

**Figure 4.**
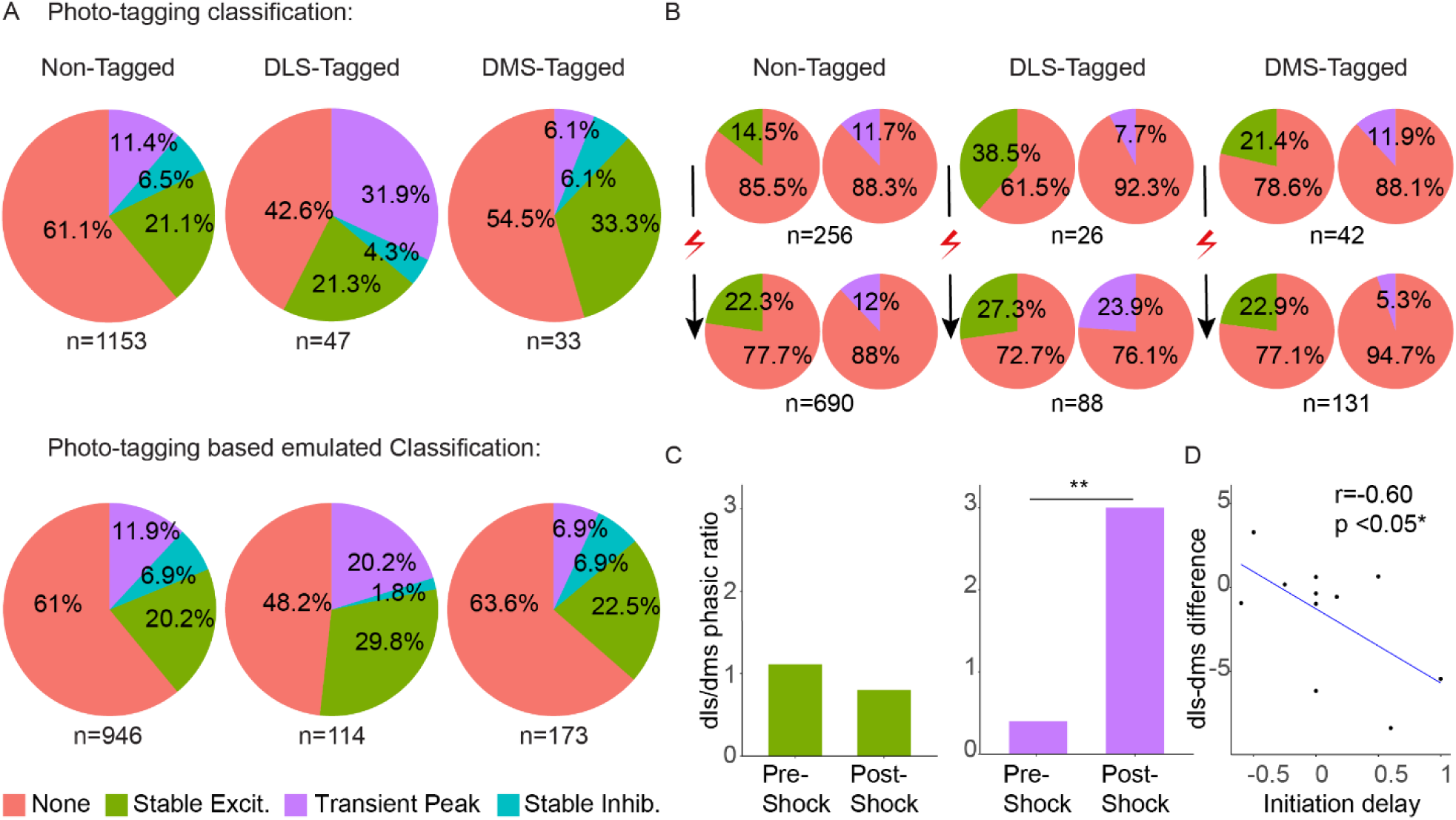
Phasic shift toward DLS following shock during grooming initiation. **(A) Top:** Distribution of the different grooming response type units in DLS-projecting, DMS-projecting and non-tagged units across all measurements, showing increased transient peak responses in the DLS projecting DA units. This pattern is replicated in the emulated classification (**bottom**), again showing enhanced transient peak responses in emulated DLS projecting DA units.**(B)** Distribution of the transient peak and stable excitation grooming response type before and after the shock, showing increased phasic activity in the DLS and decreased phasic activity in the DMS following the shock, indicating a phasic shift to the DLS.**(C)** Permutation test on the emulated classification data allowing for within-animal comparison, of the DLS/DMS unit ratio before and after the shock. No significant difference is found in stable excitation unit ratios, while a significant increase is observed in the transient peak unit ratio post-shock, indicating a transfer of phasic DA modulation to the DLS. **(D)** Regression line on a scatter plot showing the correlation between the within-animal difference in DLS phasic units and DMS phasic units, and the grooming initiation delay after the shock. A significant correlation is found, indicating that the reduction in initiation delay is associated with the transfer of phasic DA modulation to the DLS. Asterisks indicate significant comparisons (*, p<.05; **, p<.01; ***, p<.001). Pound signs indicate non-significant trends (#, .05<p<.1).

Emulating the classification using the electrophysiological properties of photo-tagged units revealed a similar result. A chi-square test revealed a significant difference in the proportion of transient peak units across groups (χ²(2) = 11.42, *p* < 0.01). Post hoc comparisons with Holm adjustment showed higher proportions of these units among the emulated DLS-projecting DA units compared to non-tagged control (*p* =0.020, adjusted *p* = 0.039), while emulated DMS-projecting DA units did not differ significantly (*p* = 0.073). A similar proportional analysis did not reveal significant differences for either stable excitation (χ²(2) = 5.77, *p* = 0.056) or stable inhibition units (χ²(2) = 4.54, *p* = 0.104; Figure 4A bottom). This similarity in the proportions firing pattern types between photo-tagged and photo-tagged emulated classification of DA units is another support for the emulated classification.

The transient peak firing pattern type seemed to stand out in DLS projecting DA units, as we already showed that the response of this unit type to grooming termination correlates with the shock effect on grooming length bout, we wanted to see whether this exclusive proportion is affected by the shock manipulation. We therefore examined the proportions of transient peak response type before and after the shock exposure (Figure5B) in - emulated by photo-tag (DLS/DMS) and non-tagged neuronal populations. To compare with potential change in other pattern types we also looked at stable excitation firing pattern type (stable inhibition type didn’t yield enough units to enable comparing proportions). Non-tagged units showed an increase only in the proportions of stable excitation response type units from 14.45% pre-shock to 22.32% post shock. A Chi-squared goodness-of-fit test indicated a significant difference between the baseline and shock conditions (X²(1) =34.53, p<0.0001). No difference was found in the proportions of transient peak responders (pre-shock 11.72% to 12.03% post shock; X²(1) =0.0642, p=0.8). There was, however, a significant increase in the proportion of transient peak firing pattern-type units from pre-to post-shock from 7.7% to 23.9% only in DLS projecting DA neurons. A Chi-squared goodness of fit test revealed a significant difference (X²(1) = 32.41, p < 0.0001). The same population of neurons showed a significant decrease in the proportion of stable excitation firing pattern-type units from 38.5% to 27.3%. A chi-squared goodness of fit test revealed a significant difference (X²(1) = 4.6545, p < 0.05). However, in DMS projecting DA neurons there was an opposite effect with a significant decrease in the proportion of transient peak firing pattern-type units from pre-to post-shock from 11.9% to 5.3%. A chi-squared test showed a significant difference (X²(1) = 5.38, p < 0.05) and no effect on the proportions of stable excitation type (X²(1) = 0.16863, p = 0.6813).

The opposite change in the proportion of transient peak units in DMS projecting DA neurons vs DLS projecting neurons correspond with the competing action control mechanisms involving the DMS and the DLS (A-O vs S-R respectively) ^36,37^ and could possibly indicate a shift in behavioral control. To test for the opposite shift, we compared the ratio of proportions of the different responding unit types after adversity in DMS and DLS projecting DA neurons and indeed found that that specifically for transient peak responding units this ratio was significantly changed towards DLS projecting DA units (Figure 4C). We conducted a non-parametric permutation test. Using the proportions of responding units in both groups, we calculated the pre-shock ratio 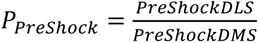, and the Post-shock ratio 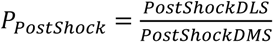. The test statistic was defined as the ratio of both ratios: 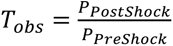 and compared to a distribution of a 10k randomly permuted test statistics. The permutation test found the post/pre shock ratios of DLS/DMS ratio to be significantly higher than the permutated population (p<0.05) only for transient peak stable excitation units. Similar permutation analysis for the ratio of stable excitation units revealed no significant difference (p=0.79). Transient peak response of DA neurons which relates to the phasic activity characterizing DA neurons ^47^. As transient peak activity was shown to shifts toward the DLS following the shock, we tested whether the observed phasic shift correlates with the behavioral effects in the post-shock measurements. As the emulated classification utilized electrophysiological properties of the units rather than original phototagging conducted in separate mice for DLS and DMS projections, we could calculate the shift of phasic control of DLS vs DMS for each animal. We found that DLS-DMS difference was inertly correlated with the initiation delay, that is the more the proportion of post-shock transient peak units tended towards the DLS the shorter it took the mouse to start grooming after grooming stimulus (Figure 4D). Correlational analysis between the DLS-DMS difference and grooming initiation delay revealed significant negative relationship (r = −0.60, p=0.05). This shift however had no detectable correlation with the duration of the shock-induced grooming bout (r = 0.21, p = 0.53) or the spontaneous )ITI( grooming duration (r < 0.001, p = 0.996).

### Activating or inhibiting DLS projecting SNc DA neurons during grooming initiation rescues grooming exacerbation after adversity

Our findings suggest that the phasic-like grooming-related transient peak DA neurons are specifically relevant for the effects of adversity on grooming behavior and that this type of response mostly characterizes SNc DA neurons projecting to the DLS. Moreover, the changes in grooming behavior following the shock manipulation are accompanied by increased ratio of these neurons. To directly test the effect of this activity on grooming, we optogenetically activated or inhibited SNc DA neurons projecting to the DLS in the same mouse during grooming induction and tested for the effects on the onset and the length of grooming both before and after the shock manipulation. To do this we injected retrograde BiPOLS^41^ to the DLS of mice and implanted optic fibers, bilaterally above the SNc (Figure 5A). Each trial, we either did not stimulate (for control trials) or intermittently light-stimulated with either blue (inhibition) or red (excitation) light, together with the grooming-inducing drop (Figure 5B).

**Figure 5.**
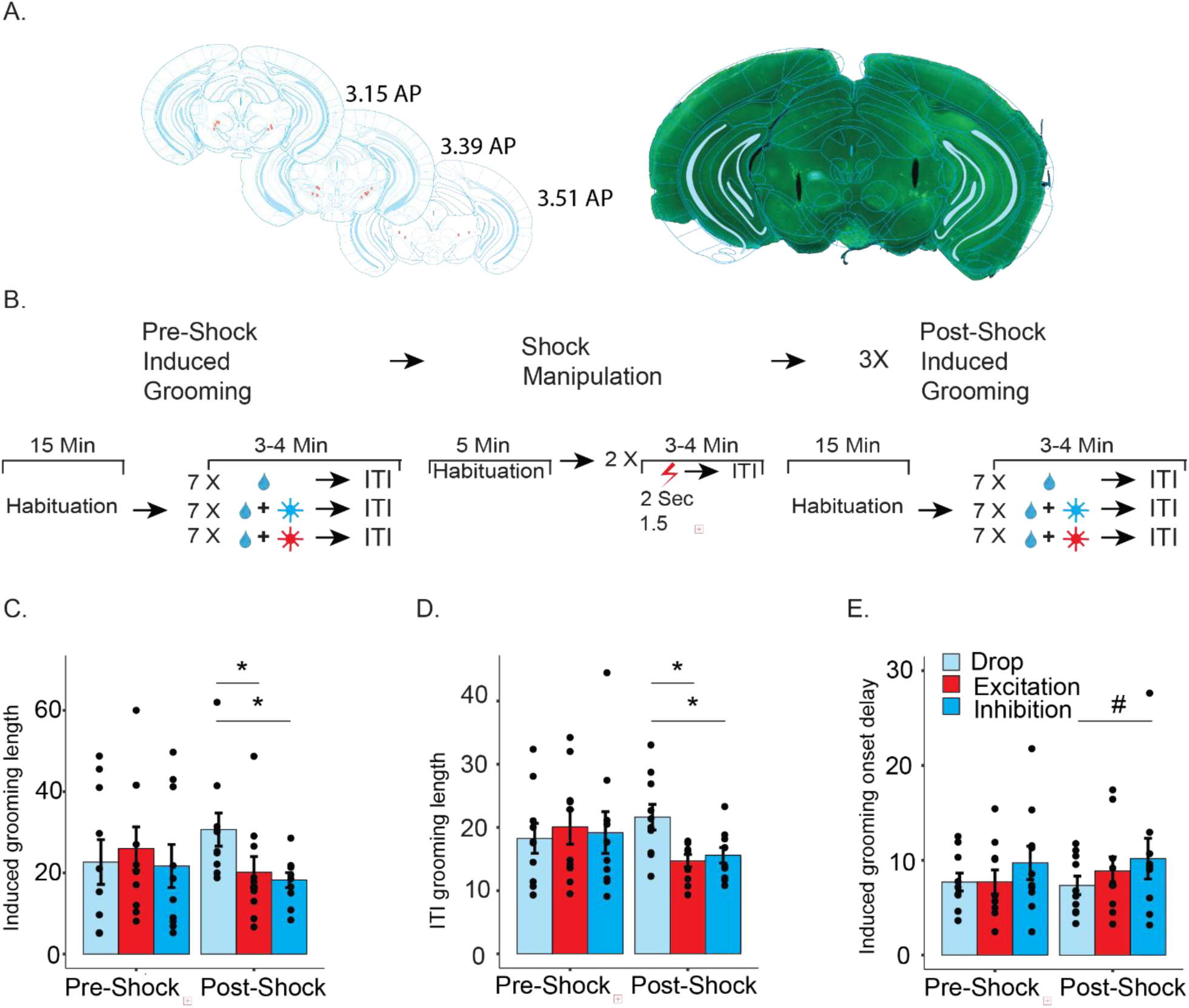
Changes in grooming behavior following shock exposure. **(A)** Left: anatomical target schemes showing the bilateral optic fibers implanted in the SNc. X’s mark histologically confirmed fiber positions from all animals included in the experiment. Right: representative coronal section showing the fibers position in SNc **(B)** Illustration of the behavioral paradigms used to assess grooming behavior combined with the stimulation, inhibition and only drop catch trials before and after the shock treatment. **(C)** Duration of induced grooming bouts in only drop catch trials as compared to the drop plus stimulation and drop plus inhibition trials. **(D)** Duration of ITI grooming bouts in only drop catch trials as compared to the drop plus stimulation and drop plus inhibition trials. **(E)** Delay in initiation of induced grooming bouts in only drop catch trials as compared to the drop plus stimulation and drop plus inhibition trials. In each plot, bars represent means of corresponding measures with error bars showing ±SEM. Asterisks indicate significant post-hoc comparisons (*, p<.05; **, p<.01; ***, p<.001). Pound signs indicate non-significant trends (#, .05<p<.1).

We found that both activation and inhibition of SNc DA neurons projecting to the DLS, immediately with the grooming induction, reduced the length of both the induced grooming immediately following the stimulation, and grooming bout much later, during the ITI. In both cases, only in measurements conducted following shock manipulation (Figures 5C-D).

When examining the length of induced bout - a two-way repeated measures ANOVA revealed no significant main effects for stimulation type (F(2, 18) = 0.97, p = 0.40) or measurement (F(1, 9) = 0,04, p = 0.84). But with a significant interaction effect (F(2, 18) = 4.67, p < 0.03). Post-hoc Dunnett’s tests revealed that both inhibition (M Difference = −12.1 seconds, p < 0.01) and excitation (M Difference = −10.7 seconds, p < 0.05) trials showed a **decrease in the length of the induced bout** compared to the water-only trials. No significant differences between either excitation or inhibition to the control group in the baseline, but we found that in the shock group (Figures 5C). Similarly in spontaneous ITI bouts, a two-way repeated measures ANOVA revealed no significant main effects for stimulation type (F(2, 18) = 2.03, p = 0.16) or for measurement (F(1, 9) = 0.92, p = 0.36), yet again revealed a significant interaction effect (F(2, 18) = 3.54, p = 0.05) with a Post-hoc Dunnett’s revealing that both inhibition (M Difference = −6, p < 0.05) and excitation (M Difference = −6.9 seconds, p < 0.01) **reduced the length of the ITI bouts** (Figure 5D). This indicated that the role of nigro-dorso striatal DA became more relevant after adversity and that the optogenetic manipulation had a lasting effect on top on its immediate effect. No effect was however, found for either stimulation or inhibition of the nigro-dorso striatal DA on the timing of the induced grooming onset, although we did see some statistical trend for the inhibition as prolonging the delay, again only after the shock manipulation (Figures 5E). A two-way repeated measures ANOVA revealed no significant main effects for stimulation type (F(2, 18) = 2.24, p = 0.14), or measurement (F(1, 9) = 0.06, p = 0.81), or stimulation type and measurement interaction (F(2, 18) = 0.17, p = 0.84). However, as we observed some mean differences between the different stimulation types, we have conducted Dunnett’s tests comparing the different stimulation type trials within each condition. In the post-shock condition, a trend was found for longer initiation delay in inhibition trials compared to no-stimulation trials (M Difference = 2.84 Seconds, p = 0.09).

## Discussion

Throughout this research, we tracked the long-term effects of anxiety-inducing, adverse electric-shock experience on physiology and grooming behavior of mice. The adverse experience exacerbated grooming in two distinct manners: It first shortened response latency to an induced grooming stimulus and second, it increased duration of grooming. The extension of grooming duration was of both the externally induced and of the spontaneously occurring grooming bouts. These findings are in line with studies who found alteration in the form of grooming and grooming exacerbation due to anxiety ^11,13,48^, and other studies who found increased reactivity ^32,38^ in animals after experiencing adverse electric-shocks.

Neuronal activity recorded in the SNc during behavior indicated three main response pattern types to grooming event: Stable-excitation units, maintaining an increase in firing rate after grooming initiation until a decrease after grooming termination. Stable inhibition units, maintaining a decrease in firing rate after grooming initiation until an increase after grooming termination. Finally-transient peak units which responded with a phasic increase in firing rate to grooming initiation and either returned to baseline or peaked again at grooming termination. Stable excitation and transient peak type units also expressed long-term changes in their grooming response following the anxiety inducing shock manipulation. Stable excitation neurons showed increased excitation in their grooming initiation response and reduced inhibition in their response to grooming termination following adversity. Transient peak units tended to peak earlier prior to the motor response as compared to pre-shock and the peak following grooming termination was increased following adversity. We also observed increased peak responses in neurons responding to grooming termination, the termination peak size also positively correlated with the duration of induced or spontaneous grooming bout. The firing pattern and the abrupt increase in firing rate of the transient peak type response units could be characterized as exhibiting phasic activity, which is a hallmark of DA signaling ^47,49^. Phasic DA release in the striatum has been shown to coincide with the onset and offset of action sequences, suggesting a crucial role in action initiation and termination ^50–52^. The earlier initiation peak and stronger termination peak which correlate with the prolonging of grooming following adversity suggest these transient, phasic response of grooming related SNc neurons play a role in anxiety related changes in grooming.

Using photo-tagging in DAT-Cre mice we managed to identify these response type neurons as characterizing mostly SNc DA neurons projecting to the DLS. Moreover, we observed an increase in their proportion in SNc DA neurons projecting to the DLS following anxiety induction, while SNc DA neurons projecting to the DMS showed a decreased proportion of the same type of neurons. In fact, the DLS/DMS projecting proportion difference of phasic responders SNc DA neurons had a strong negative correlation with grooming initiation delay, characterizing the post-shock grooming response. This DLS/DMS ratio corresponds with the well-documented competitive interaction between action control systems – the S-R (regulated by DLS activity) and A-O (regulated by DMS activity) control ^53–56^. The shift of behavioral change relevant DA neurons to a more prevalent input into the DLS over the DMS following adversity, can indicate an anxiety related shift in action control towards S-R dominated control ^57–59^.

The result of activating or inhibiting DLS-projecting SNc DA neurons, during the approximated time of phasic response, at grooming induction, further underscores the role these neurons play in the effect of anxiety on grooming. Interestingly either exciting or inhibiting those neurons, reduced most of the anxiety-induced grooming effects. An effect that was selective only to measurements which followed the shock manipulation, suggesting that the activity of DLS projecting DA neurons becomes more critical for grooming regulation in an anxious state.

The observation that both inhibition and excitation of DLS-projecting DA neurons similarly reduced the duration of DA-related grooming, indicates that the DA signal might be precisely synchronized with the grooming, as could also be inferred from the grooming dependent response types of DA neurons recorded in the SNc. Therefore, either a fixed rate excitation train or inhibition at sensitive timing can disrupt the DA signal and interrupt the role of these neurons in executing the planned action.

Stress was indeed suggested to bias circuit dynamics toward S-R neural circuits dominance ^57–59^, which supports the exacerbation of grooming through a change in DLS-DMS equilibrium. Joining previous findings, showing that the adjusting of grooming behavior by midbrain DA activity, differs based on the target area of these DA neurons ^21,24^, the results of this work provide novel evidence for the specific role of DLS projecting DA phasic neurons in this control system shift.

The exacerbation of repetitive behaviors under stress is central to conditions like OCD, which is suggested as an imbalance between habitual and goal directed behavior ^60–64^. The same imbalance that was characterized in excessive grooming rodent models of OCD ^65,66^. Our findings provide evidence for different involvement of DA neurons in onset, duration and termination of grooming, and identifies specific DA neuronal type which specifically targets the DLS involved in increased intensiveness of grooming following adversity.

## Acknowledgements

We thank all Klavir lab members for discussions and critical reading of the manuscript.

## Author contribution

O.K. directed the study. Experimental design, L.G and O.K, data acquisition, and interpretation, L.G, L.A, S.E and O.K.; manuscript writing, L.G., and O.K.

## References

1. Lüthi, A. & Lüscher, C. Pathological circuit function underlying addiction and anxiety disorders. Nat Neurosci 17, 1635–1643 (2014).

2. Milad, M. R. & Rauch, S. L. Obsessive-compulsive disorder: beyond segregated cortico-striatal pathways. Trends Cogn Sci 16, 43–51 (2012).

3. Adams, T. G. et al. The Role of Stress in the Pathogenesis and Maintenance of Obsessive-Compulsive Disorder. Chronic Stress 2, 2470547018758043 (2018).

4. Baribeau, D. A. et al. Repetitive Behavior Severity as an Early Indicator of Risk for Elevated Anxiety Symptoms in Autism Spectrum Disorder. Journal of the American Academy of Child & Adolescent Psychiatry 59, 890–899.e3 (2020).

5. Buse, J., Kirschbaum, C., Leckman, J. F., Münchau, A. & Roessner, V. The Modulating Role of Stress in the Onset and Course of Tourette’s Syndrome: A Review. Behav Modif 38, 184– 216 (2014).

6. Bienvenu, O. J., et al. *Sapap3* and pathological grooming in humans: Results from the OCD collaborative genetics study. American J of Med Genetics Pt B **150B**, 710–720 (2009).

7. Feusner, J. D., Hembacher, E. & Phillips, K. A. The Mouse Who Couldn’t Stop Washing: *Pathologic Grooming in Animals and Humans*. CNS spectr. 14, 503–513 (2009).

8. Kalueff, A. V. et al. Neurobiology of rodent self-grooming and its value for translational neuroscience. Nat Rev Neurosci 17, 45–59 (2016).

9. Garner, J. P. et al. Reverse-translational biomarker validation of Abnormal Repetitive Behaviors in mice: an illustration of the 4P’s modeling approach. Behav Brain Res 219, 189– 196 (2011).

10. Kalueff, A. V., Aldridge, J. W., LaPorte, J. L., Murphy, D. L. & Tuohimaa, P. Analyzing grooming microstructure in neurobehavioral experiments. Nat Protoc 2, 2538–2544 (2007).

11. Kalueff, A. V. & Tuohimaa, P. Mouse grooming microstructure is a reliable anxiety marker bidirectionally sensitive to GABAergic drugs. Eur J Pharmacol 508, 147–153 (2005).

12. Pires, G. N., Tufik, S. & Andersen, M. L. Grooming analysis algorithm: Use in the relationship between sleep deprivation and anxiety-like behavior. Progress in Neuro-Psychopharmacology and Biological Psychiatry 41, 6–10 (2013).

13. Smolinsky, A. N., Bergner, C. L., LaPorte, J. L. & Kalueff, A. V. Analysis of Grooming Behavior and Its Utility in Studying Animal Stress, Anxiety, and Depression. in Mood and Anxiety Related Phenotypes in Mice (ed. Gould, T. D.) vol. 42 21–36 (Humana Press, Totowa, NJ, 2009).

14. Liu, H. et al. Dissection of the relationship between anxiety and stereotyped self-grooming using the Shank3B mutant autistic model, acute stress model and chronic pain model. Neurobiology of Stress 15, 100417 (2021).

15. Wang, B. et al. Zfp462 deficiency causes anxiety-like behaviors with excessive self-grooming in mice. Genes Brain and Behavior 16, 296–307 (2017).

16. Denys, D., Zohar, J. & Westenberg, H. G. M. The role of dopamine in obsessive-compulsive disorder: preclinical and clinical evidence. J Clin Psychiatry 65 **Suppl 14**, 11–17 (2004).

17. Peters, S. K., Dunlop, K. & Downar, J. Cortico-Striatal-Thalamic Loop Circuits of the Salience Network: A Central Pathway in Psychiatric Disease and Treatment. Front Syst Neurosci 10, 104 (2016).

18. Casado-Sainz, A. et al. Dorsal striatal dopamine induces fronto-cortical hypoactivity and attenuates anxiety and compulsive behaviors in rats. Neuropsychopharmacol. 47, 454–464 (2022).

19. Seiler, J. L. et al. Dopamine signaling in the dorsomedial striatum promotes compulsive behavior. Current Biology 32, 1175–1188.e5 (2022).

20. Szechtman, H., Sulis, W. & Eilam, D. Quinpirole induces compulsive checking behavior in rats: A potential animal model of obsessive-compulsive disorder (OCD). Behavioral Neuroscience 112, 1475–1485 (1998).

21. Xue, J. et al. Midbrain dopamine neurons arbiter OCD-like behavior. Proc Natl Acad Sci U S A 119, e2207545119 (2022).

22. Wood, J., LaPalombara, Z. & Ahmari, S. E. Monoamine abnormalities in the SAPAP3 knockout model of obsessive-compulsive disorder-related behaviour. Phil. Trans. R. Soc. B 373, 20170023 (2018).

23. Manning, E. E., Wang, A. Y., Saikali, L. M., Winner, A. S. & Ahmari, S. E. Disruption of prepulse inhibition is associated with compulsive behavior severity and nucleus accumbens dopamine receptor changes in Sapap3 knockout mice. Sci Rep 11, 9442 (2021).

24. Givon, L., Edut, S. & Klavir, O. The role of fear and dopamine-striatal pathways in grooming. Neuropharmacology 269, 110323 (2025).

25. Lykouras, L., Alevizos, B., Michalopoulou, P. & Rabavilas, A. Obsessive–compulsive symptoms induced by atypical antipsychotics. A review of the reported cases. Progress in Neuro-Psychopharmacology and Biological Psychiatry 27, 333–346 (2003).

26. Sareen, J. et al. Do antipsychotics ameliorate or exacerbate Obsessive Compulsive Disorder symptoms? Journal of Affective Disorders 82, 167–174 (2004).

27. Crum, R. M. & Anthony, J. C. Cocaine use and other suspected risk factors for obsessive-compulsive disorder: a prospective study with data from the Epidemiologic Catchment Area surveys. Drug and Alcohol Dependence 31, 281–295 (1993).

28. King, J., Dowling, N. & Leow, F. Methylphenidate in the treatment of an adolescent female with obsessive-compulsive disorder and attention deficit hyperactivity disorder: a case report. Australas Psychiatry 25, 178–180 (2017).

29. Koran, L. M., Aboujaoude, E. & Gamel, N. N. Double-blind study of dextroamphetamine versus caffeine augmentation for treatment-resistant obsessive-compulsive disorder. J Clin Psychiatry 70, 1530–1535 (2009).

30. Vulink, N. C. C., Denys, D. & Westenberg, H. G. M. Bupropion for Patients With Obsessive-Compulsive Disorder: An Open-Label, Fixed-Dose Study. J. Clin. Psychiatry 66, 228–230 (2005).

31. Cai, H. et al. Amygdalo-nigral circuit mediates stress-induced vulnerability to the parkinsonian toxin MPTP. CNS Neurosci Ther 29, 1940–1952 (2023).

32. Yao, L. et al. Stress Controllability Modulates Basal Activity of Dopamine Neurons in the Substantia Nigra Compacta. eNeuro 8, ENEURO.0044-21.2021 (2021).

33. Haber, S. N., Fudge, J. L. & McFarland, N. R. Striatonigrostriatal pathways in primates form an ascending spiral from the shell to the dorsolateral striatum. J Neurosci 20, 2369–2382 (2000).

34. Basile, G. A. et al. Striatal topographical organization: Bridging the gap between molecules, connectivity and behavior. Eur J Histochem 65, 3284 (2021).

35. Menegas, W. et al. Dopamine neurons projecting to the posterior striatum form an anatomically distinct subclass. Elife 4, e10032 (2015).

36. Redgrave, P. et al. Goal-directed and habitual control in the basal ganglia: implications for Parkinson’s disease. Nat Rev Neurosci 11, 760–772 (2010).

37. Yin, H. H. & Knowlton, B. J. The role of the basal ganglia in habit formation. Nat Rev Neurosci 7, 464–476 (2006).

38. Burguière, E., Monteiro, P., Feng, G. & Graybiel, A. M. Optogenetic Stimulation of Lateral Orbitofronto-Striatal Pathway Suppresses Compulsive Behaviors. Science 340, 1243–1246 (2013).

39. Ritter, A., Habusha, S., Givon, L., Edut, S. & Klavir, O. Prefrontal control of superior colliculus modulates innate escape behavior following adversity. Nat Commun 15, 2158 (2024).

40. Siegmund, A. & Wotjak, C. T. Toward an Animal Model of Posttraumatic Stress Disorder. Annals of the New York Academy of Sciences 1071, 324–334 (2006).

41. Simon, P., Dupuis, R. & Costentin, J. Thigmotaxis as an index of anxiety in mice. Influence of dopaminergic transmissions. Behav Brain Res 61, 59–64 (1994).

42. Vierock, J. et al. BiPOLES is an optogenetic tool developed for bidirectional dual-color control of neurons. Nat Commun 12, 4527 (2021).

43. Mathis, A. et al. DeepLabCut: markerless pose estimation of user-defined body parts with deep learning. Nat Neurosci 21, 1281–1289 (2018).

44. Ungless, M. A. & Grace, A. A. Are you or aren’t you? Challenges associated with physiologically identifying dopamine neurons. Trends in Neurosciences 35, 422–430 (2012).

45. Chinta, S. J. & Andersen, J. K. Dopaminergic neurons. The International Journal of Biochemistry & Cell Biology 37, 942–946 (2005).

46. Nair-Roberts, R. G. et al. Stereological estimates of dopaminergic, GABAergic and glutamatergic neurons in the ventral tegmental area, substantia nigra and retrorubral field in the rat. Neuroscience 152, 1024–1031 (2008).

47. Lerner, T. N. et al. Intact-Brain Analyses Reveal Distinct Information Carried by SNc Dopamine Subcircuits. Cell 162, 635–647 (2015).

48. Schultz, W., Dayan, P. & Montague, P. R. A Neural Substrate of Prediction and Reward. Science 275, 1593–1599 (1997).

49. Granjeiro, É. M. et al. Behavioral and Cardiorespiratory Responses to Bilateral Microinjections of Oxytocin into the Central Nucleus of Amygdala of Wistar Rats, an Experimental Model of Compulsion. PLoS ONE 9, e99284 (2014).

50. Grace, A. A., Floresco, S. B., Goto, Y. & Lodge, D. J. Regulation of firing of dopaminergic neurons and control of goal-directed behaviors. Trends in Neurosciences 30, 220–227 (2007).

51. Jin, X. & Costa, R. M. Start/stop signals emerge in nigrostriatal circuits during sequence learning. Nature 466, 457–462 (2010).

52. Ko, D. & Wanat, M. J. Phasic Dopamine Transmission Reflects Initiation Vigor and Exerted Effort in an Action- and Region-Specific Manner. J Neurosci 36, 2202–2211 (2016).

53. Wassum, K. M., Ostlund, S. B. & Maidment, N. T. Phasic Mesolimbic Dopamine Signaling Precedes and Predicts Performance of a Self-Initiated Action Sequence Task. Biological Psychiatry 71, 846–854 (2012).

54. Bergstrom, H. C. et al. Dorsolateral Striatum Engagement Interferes with Early Discrimination Learning. Cell Reports 23, 2264–2272 (2018).

55. Dezfouli, A. & Balleine, B. W. Actions, Action Sequences and Habits: Evidence That Goal-Directed and Habitual Action Control Are Hierarchically Organized. PLoS Comput Biol 9, e1003364 (2013).

56. Hardwick, R. M., Forrence, A. D., Krakauer, J. W. & Haith, A. M. Time-dependent competition between goal-directed and habitual response preparation. Nat Hum Behav 3, 1252–1262 (2019).

57. Turner, K. M., Svegborn, A., Langguth, M., McKenzie, C. & Robbins, T. W. Opposing Roles of the Dorsolateral and Dorsomedial Striatum in the Acquisition of Skilled Action Sequencing in Rats. J. Neurosci. 42, 2039–2051 (2022).

58. Giovanniello, J. R. et al. A dual-pathway architecture for stress to disrupt agency and promote habit. Nature 640, 722–731 (2025).

59. Schwabe, L. & Wolf, O. T. Stress Prompts Habit Behavior in Humans. J. Neurosci. 29, 7191–7198 (2009).

60. Wirz, L., Bogdanov, M. & Schwabe, L. Habits under stress: mechanistic insights across different types of learning. Current Opinion in Behavioral Sciences 20, 9–16 (2018).

61. Banca, P. et al. Imbalance in habitual versus goal directed neural systems during symptom provocation in obsessive-compulsive disorder. Brain 138, 798–811 (2015).

62. Gillan, C. M. et al. Disruption in the Balance Between Goal-Directed Behavior and Habit Learning in Obsessive-Compulsive Disorder. AJP 168, 718–726 (2011).

63. Simmler, L. D. & Ozawa, T. Neural circuits in goal-directed and habitual behavior: Implications for circuit dysfunction in obsessive-compulsive disorder. Neurochemistry International 129, 104464 (2019).

64. Voon, V. et al. Disorders of compulsivity: a common bias towards learning habits. Mol Psychiatry 20, 345–352 (2015).

65. Xu, C. et al. Imbalance in functional and structural connectivity underlying goal-directed and habitual learning systems in obsessive-compulsive disorder. Cerebral Cortex 32, 3690–3705 (2022).

66. Ehmer, I. et al. Evidence for Distinct Forms of Compulsivity in the SAPAP3 Mutant-Mouse Model for Obsessive-Compulsive Disorder. eNeuro 7, ENEURO.0245-19.2020 (2020).

67. Hadjas, L. C., Lüscher, C. & Simmler, L. D. Aberrant habit formation in the Sapap3-knockout mouse model of obsessive-compulsive disorder. Sci Rep 9, (2019).

