## Supplementary material for "Post Adversity Changes in Nigro-Striatal Dopamine: A Mechanism for Anxiety Induced Exacerbated Innate Repetitive Behaviors"

### Supplementary figures:

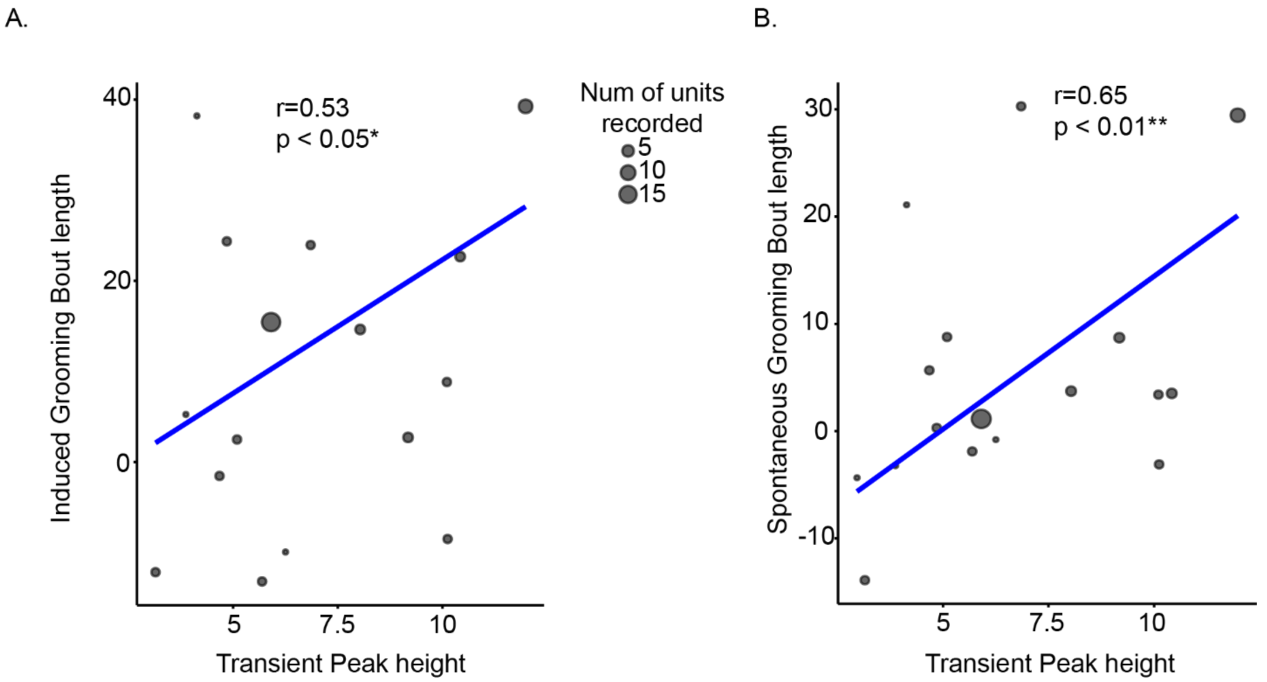


**Supplementary Figure 1. Correlation between grooming length and transient peak responses to grooming termination.**

**(A)** A regression line over a scatter plot showing a weighted regression between the normalized length of induced grooming bouts and the average response magnitude (peak height) of transient peak units responding to grooming termination indicating that higher termination peak responses are associated with longer grooming bouts. The number of transient peak units per measurement was used as the weighting factor. **(B)** A regression line over a scatter plot showing a weighted regression between the normalized average length of inter-trial interval (ITI) bouts and the average response magnitude (peak height) of transient peak units responding to grooming termination suggesting that termination peak magnitude also scales with the duration of preceding ITI bouts. The number of transient peak units per measurement was again used as the weighting factor. Asterisks indicate significant correlation (*, p<.05; **, p<.01; ***, p<.001).

### Supplementary tables:

### Full statistics tables by figures:

| Figure | Dependent Variable | Group | N | Mean | SD |
|---|---|---|---|---|---|
| 1B | Grooming Initiation Delay | Control Measure1 | 9 | 0.66 | 5.18 |
| | | Control Measure2 | 9 | 0.23 | 1.77 |
| | | Control Measure3 | 9 | 1.91 | 4.24 |
| | | Shock Measure1 | 17 | -1.22 | 2.52 |
| | | Shock Measure2 | 17 | -0.71 | 3.97 |
| | | Shock Measure3 | 17 | -1.19 | 4.1 |
| | | **Effect** | **Test** | **Statistic** | **p** |
| | | Group Variances Comparison | F-Test | F(50,26)=0.82 | p = 0.54 |
| | | Group | Mixed-ANOVA | F(1,25)=2.97 | p=0.098 |
| | | Measure | Mixed-ANOVA | F(2,48)=1.4 | p=0.88 |
| | | Group x Measure | Mixed-ANOVA | F(2,48)=0.74 | p=0.48 |
| | | **Group** | **N** | **Mean** | **SD** |
| 1B - small | Grooming Initiation Delay | Control | 28 | 1.39 | 2.75 |
| | | Shock | 12 | -1.39 | 3.90 |
| | | **Effect** | **Test** | **Statistic** | **p** |
| | | Group | F-Test | F(27,11)=2.02 | p=0.22 |
| | | Group | T-test | t(38)=2.24 | P = 0.031 |
| **Figure** | **Dependent Variable** | **Group** | **N** | **Mean** | **SD** |
| 1C | Grooming Initiation Length | Control Measure1 | 11 | -1.89 | 7.84 |
| | | Control Measure2 | 11 | 2.98 | 5.67 |
| | | Control Measure3 | 11 | -1.82 | 5.90 |
| | | Shock Measure1 | 19 | 0.64 | 9.08 |
| | | Shock Measure2 | 19 | 12.07 | 15.32 |
| | | Shock Measure3 | 19 | 7.78 | 8.67 |

| | | | | | |
|---|---|---|---|---|---|
| | | Effect | Test | Statistic | p |
| | | Group Variances Comparison | F-Test | F(56,32)=3.27 | p= 0.0005 |
| | | Group | Mixed-Type III Anova with Welch Correction | F(1,28)=6.34 | p=0.018 |
| | | Measure | Mixed-Type III Anova with Welch Correction | F(1.81,50.77)=6.98 | p=0.003 |
| | | Group x Measure | Mixed-Type III Anova with Welch Correction | F(1.81,50.77)=1.62 | p=0.21 |
| | | Group | N | Mean | SD |
| 1C small | Grooming Initiation Length | Control | 12 | 0.309 | 5.91 |
| | | Shock | 28 | 5.70 | 10.7 |
| | | Effect | Test | Statistic | p |
| | | Group | F-Test | F(27,11)=3.28 | p=0.042 |
| | | Group | T-test | T(35.28)=-2.04 | p=0.049 |
| Figure | Dependent Variable | Group | N | Mean | SD |
| 1D | Grooming ITI Length | Control Measure1 | 11 | -1.19 | 2.57 |
| | | Control Measure2 | 11 | -1.15 | 4.08 |
| | | Control Measure3 | 11 | -1.31 | 3.63 |
| | | Shock Measure1 | 20 | 3.6 | 4.5 |
| | | Shock Measure2 | 20 | 5.94 | 10.44 |
| | | Shock Measure3 | 20 | 4.82 | 8.54 |
| | | Effect | Test | Statistic | p |
| | | Group Variances Comparison | F-Test | F(59,32)=6.24 | p= 0.0000003 |
| | | Group | Mixed-Type III Anova | F(1,29)=5.71 | p=0.024 |

| | | | | | |
|---|---|---|---|---|---|
| | | | with Welch Correction | | |
| | | Measure | Mixed-Type III Anova with Welch Correction | F(51.31,31.02)=2.25 | p=0.122 |
| | | Group x Measure | Mixed-Type III Anova with Welch Correction | F(51.31,31.02)=2.27 | p=0.119 |
| | | Group | N | Mean | SD |
| 1D small | Grooming ITI Length | Control | 12 | -1.32 | 2.19 |
| | | Shock | 28 | 3.01 | 6.02 |
| | | Effect | Test | Statistic | p |
| | | Group | F-Test | F(27,11)=7.54 | p=0.0012 |
| | | Group | T-test | t(37.49)=-3.32 | P=0.0019 |
| Figure | Dependent Variable | Group | N | Mean | SD |
| 1E | OpenField Center Distance | Control Measure1 | 8 | -77.87 | 41.68 |
| | | Control Measure2 | 8 | -80.42 | 96.22 |
| | | Control Measure3 | 8 | -59.11 | 62.54 |
| | | Shock Measure1 | 9 | 1.65 | 75.27 |
| | | Shock Measure2 | 9 | 72.27 | 169.39 |
| | | Shock Measure3 | 9 | 58.97 | 126 |
| | | Effect | Test | Statistic | p |
| | | Group Variances Comparison | F-Test | F(26,23)=3.55 | p= 0.0031 |
| | | Group | Mixed-Type III Anova with Welch Correction | F(1,15.06)=6.46 | p=0.023 |
| | | Measure | Mixed-Type III Anova with Welch Correction | F(1,54,23.06)=2.14 | p=0.149 |

| | | | | | |
|---|---|---|---|---|---|
| | | Group x Measure | Mixed-Type III Anova with Welch Correction | F(1,54,23.06)=1.64 | P=0.217 |
| | | Group | N | Mean | SD |
| 1E small | Openfield Center Distance | Control | 10 | -73.8 | 65.7 |
| | | Shock | 12 | 28.4 | 118 |
| | | Effect | Test | Statistic | p |
| | | Group | F-Test | F(9,11)=0.31 | p=0.09 |
| | | Group | T-test | t(20)=-2.44 | P=0.0239 |

| Figure | Dependent Variable | Group | N | Mean | SD |
|---|---|---|---|---|---|
| 1F | OpenField Center Time % | Control Measure1 | 8 | -0.02 | 0.11 |
| | | Control Measure2 | 8 | 0.03 | 0.12 |
| | | Control Measure3 | 8 | 0.03 | 0.1 |
| | | Shock Measure1 | 9 | -0.07 | 0.1 |
| | | Shock Measure2 | 9 | -0.1 | 0.22 |
| | | Shock Measure3 | 9 | -0.11 | 0.13 |
| | | Effect | Test | Statistic | p |
| | | Group Variances Comparison | F-Test | F(26,23)=2.046 | p=0.086 |
| | | Group | Mixed-Anova | F(1,15)=4.27 | p = 0.056 |
| | | Measure | Mixed-Anova | F(2,30)=0.045 | p=0.956 |
| | | Group x Measure | Mixed-Anova | F(2,30)=0.796 | p=0.461 |
| | | Group | N | Mean | SD |
| 1F small | OpenField Center Time % | Control | 10 | 0.015 | 0.06 |
| | | Shock | 12 | -0.09 | 0.12 |
| | | Effect | Test | Statistic | p |
| | | Group | F-Test | F(9,11)=0.274 | p=0.06 |
| | | Group | T-test | t(20)=2.35 | p=0.029 |

| Figure | Dependent Variable | Group | N | Mean | SD |
|---|---|---|---|---|---|
| 2D | Grooming Initiation Stable Excitation Units Magnitude | Group | N | Mean | SD |
| | | Pre-Shock | 55 | 1.49 | 0.41 |
| | | Post-Shock | 202 | 3.60 | 4.36 |
| | | Effect | Test | Statistic | p |
| | | Group | F-Test | F(54,201)=0.008 | p<0.00001 |
| | | Group | T-test | t(213.27)=-6.77 | P = 0.00001 |
| 2D | Grooming Initiation Stable Excitation Units Timing | Group | N | Mean | SD |
| | | Pre-Shock | 55 | 91.6 | 124 |
| | | Post-Shock | 202 | 136 | 131 |
| | | Effect | Test | Statistic | p |
| | | Group | F-Test | F(54,201)=0.90 | p = 0.672 |
| | | Group | T-test | t(255)=-2.26 | P = 0.0249 |
| 2F | Grooming Termination Stable Inhibition Units Magnitude | Group | N | Mean | SD |
| | | Pre-Shock | 55 | 0.97 | 0.17 |
| | | Post-Shock | 202 | 1.22 | 0.58 |
| | | Effect | Test | Statistic | p |
| | | Group | F-Test | F(58,167)=0.88 | P < 0.00001 |
| | | Group | T-test | t(221.18)=-4.94 | P < 0.0001 |
| 2I | Grooming Initiation Transient Peak Units Peak Height | Group | N | Mean | SD |
| | | Pre-Shock | 59 | 8.11 | 5.37 |
| | | Post-Shock | 168 | 7.44 | 3.63 |
| | | Effect | Test | Statistic | p |
| | | Group | F-Test | F(35,105)=2.189 | P=0.024 |
| | | Group | T-test | t(46.33)=1.43 | P=0.159 |
| 2I | Grooming Initiation Transient Peak Units Timing | Group | N | Mean | SD |
| | | Pre-Shock | 59 | 94.7 | 161 |
| | | Post-Shock | 168 | 42.4 | 176 |
| | | Effect | Test | Statistic | p |
| | | Group | F-Test | F(35,105)=0.841 | P=0.57 |
| | | Group | T-Test | t(140)=1.88 | p=0.063 |
| | | Group | N | Mean | SD |

| | | | | | |
|---|---|---|---|---|---|
| 2K | Grooming Termination Transient Peak Units Peak Height | Pre-Shock | 25 | 5.98 | 2.18 |
| | | Post-Shock | 58 | 7.67 | 3.94 |
| | | Effect | Test | Statistic | p |
| | | Group | F-Test | F(24,57)=0.31 | p=0.022 |
| | | Group | T-Test | t(75.88)=-2.49 | P=0.014 |
| 2K | Grooming Termination Transient Peak Units Timing | Group | N | Mean | SD |
| | | Pre-Shock | 25 | -6.13 | 146 |
| | | Post-Shock | 58 | -5.3 | 167 |
| | | Effect | Test | Statistic | p |
| | | Group | F-Test | F(24,57)=1.04 | p=0.88 |
| | | Group | T-Test | t(81)=-0.02 | p=0.98 |
| 2N | Grooming Initiation Stable Inhibition Magnitude | Group | N | Mean | SD |
| | | Pre-Shock | 25 | 0.99 | 0.22 |
| | | Post-Shock | 54 | 1.01 | 0.26 |
| | | Effect | Test | Statistic | p |
| | | Group | F-Test | F(23,51)=0.74 | p = 0.429 |
| | | Group | T-Test | t(74)=-0.34 | P=0.74 |
| | | Group | N | Mean | SD |
| 2P | Grooming Termination Stable Excitation Magnitude | Pre-Shock | 39 | 2.01 | 0.87 |
| | | Post-Shock | 50 | 1.77 | 0.85 |
| | | Effect | Test | Statistic | p |
| | | Group | F-Test | F(49,38)=0.95 | p=0.86 |
| | | Group | T-Test | t(87)=1.32 | p=0.19 |
| 2P | Grooming Termination Stable Excitation Timing | Group | N | Mean | SD |
| | | Pre-Shock | 39 | -367 | 173 |
| | | Post-Shock | 50 | -394 | 186 |
| | | Effect | Test | Statistic | p |
| | | Group | F-Test | F(49,38-1.15) | P=0.662 |
| | | Group | T-test | t(87)=0.70 | P=0.488 |

| Figure | Dependent Variable | Group | N | Mean | SD |
|---|---|---|---|---|---|
| 3E - Top | FR | Group | N | Mean | SD |
| | | DLS | 46 | 6.40 | 3.64 |
| | | Other | 1158 | 7.44 | 6.63 |
| | | Effect | Test | Statistic | p |
| | | Group | F-Test | F(45/1157)= 0.302 | P < 0.00001 |
| | | Group | T-test | t(57.60)=-2.34 | P=0.022 |
| 3E - Right | FWHM | Group | N | Mean | SD |
| | | DLS | 47 | 110 | 29.7 |
| | | Other | 1177 | 115 | 39.2 |
| | | Effect | Test | Statistic | p |
| | | Group | F-Test | F(46,1176)= 0.575 | p = 0.019 |
| | | Group | T-test | t(52.60)=-1.16 | p=0.25 |
| 3F - Top | FR | Group | N | Mean | SD |
| | | DMS | 33 | 9.66 | 6.39 |
| | | Other | 1172 | 7.64 | 6.55 |
| | | Effect | Test | Statistic | p |
| | | Group | F-Test | F(32/1171)= 0.95 | p=0.92 |
| | | Group | T-test | t(1203)=1.75 | p=0.08 |
| 3F - Right | FWHM | Group | N | Mean | SD |
| | | DMS | 33 | 88.8 | 12.2 |
| | | Other | 1191 | 116 | 39.1 |
| | | Effect | Test | Statistic | p |
| | | Group | F-Test | F(32,1190)0.097 | P<0.00001 |
| | | Group | T-test | t(52.705)=-11.175 | P<0.00001 |
| 3G - Top | FR | Group | N | Mean | SD |
| | | DLS | 46 | 6.40 | 3.64 |
| | | DMS | 33 | 9.66 | 6.39 |
| | | Effect | Test | Statistic | p |
| | | Group | F-Test | F(45,32)=0.32 | P=0.005 |
| | | Group | T-test | t(46.80)=-2.64 | p=0.011 |

| | | Group | N | Mean | SD |
|---|---|---|---|---|---|
| 3G - Right | FWHM | DLS | 47 | 110 | 29.7 |
| | | DMS | 33 | 88.8 | 12.2 |
| | | Effect | Test | Statistic | p |
| | | Group | F-Test | F(46,32)=5.93 | P<0.00001 |
| | | Group | T-test | t(65.32)=4.38 | P=0.00004 |

| Figure | Dependent Variable | Unit Type | N |
|---|---|---|---|
| 4A Top - Left | Grooming Initiation Unit Type | Transient Peak | 131 |
| | | Stable Excitation | 243 |
| | | Stable Inhibition | 75 |
| | | None | 704 |
| 4A Top - Middle | Grooming Initiation Unit Type | Unit Type | N |
| | | Transient Peak | 15 |
| | | Stable Excitation | 10 |
| | | Stable Inhibition | 2 |
| | | None | 20 |
| 4A Top - Right | Grooming Initiation Unit Type | Unit Type | N |
| | | Transient Peak | 2 |
| | | Stable Excitation | 11 |
| | | Stable Inhibition | 2 |
| | | None | 18 |

| Figure | Effect | Test | Statistic | p |
|---|---|---|---|---|
| 4A Top | Stable Excitation | Chi-Square Test | $\chi^2(2) = 2.86$ | P=0.24 |
| | Stable Inhibition | Chi-Square Test | $\chi^2(2) = 0.39$ | P=0.82 |
| | Transient Peak | Chi-Square Test | $\chi^2(2) = 19.195$ | P=0.000068 |
| | DLS vs Non-Tagged Transient Peak | Chi-Square Test | $\chi^2(2) = 15.98$ | P=0.00012 (Adjusted with Holm Correction) |
| | DMS vs Non-Tagged Transient Peak | Chi-Square Test | $\chi^2(2) = 0.45$ | p=0.5017 (Adjusted with Holm Correction) |

| | | | |
|---|---|---|---|
| 4A Bottom – Left | Grooming Initiation Unit Type | **Unit Type** | **N** |
| | | Transient Peak | 113 |
| | | Stable Excitation | 191 |
| | | Stable Inhibition | 65 |
| | | None | 946 |
| 4A Bottom – Middle | Grooming Initiation Unit Type | **Unit Type** | **N** |
| | | Transient Peak | 23 |
| | | Stable Excitation | 34 |
| | | Stable Inhibition | 2 |
| | | None | 55 |
| 4A Bottom – Right | Grooming Initiation Unit Type | **Unit Type** | **N** |
| | | Transient Peak | 12 |
| | | Stable Excitation | 39 |
| | | Stable Inhibition | 12 |
| | | None | 110 |
| 4A – Bottom | Effect | Test | Statistic |

| | | |
|---|---|---|
| Stable Excitation (Emulated) | Chi-Square Test | $\chi^2(2) = 5.77$ |
| Stable Inhibition (Emulated) | Chi-Square Test | $\chi^2(2) = 4.5358$ |
| Transient Peak (Emulated) | Chi-Square Test | $\chi^2(2) = 11.42$ |
| DLS vs Non-Classified Transient Peak (Emulated) | Chi-Square Test | $\chi^2(2) = 5.448$ |
| DMS vs Non- Classified Transient Peak (Emulated) | Chi-Square Test | $\chi^2(2) = 3.210$ |

4B

| Classification (Emulated) | Period | Unit Type |
|---|---|---|
| Non-Classified | Pre-Shock | Transient Peak |
| | | Stable Excitation |
| | | Stable Inhibition |
| | | None |
| | Post-Shock | Transient Peak |
| | | Stable Excitation |
| | | Stable Inhibition |
| | | None |
| Classification (Emulated) | Period | Unit Type |
| DLS | Pre-Shock | Transient Peak |
| | | Stable Excitation |
| | | Stable Inhibition |
| | | None |
| | Post-Shock | Transient Peak |
| | | Stable Excitation |
| | | Stable Inhibition |
| | | None |
| Classification (Emulated) | Period | Unit Type |
| DMS | Pre-Shock | Transient Peak |
| | | Stable Excitation |

| | | | | |
|---|---|---|---|---|
| | | | Stable Inhibition | |
| | | | None | |
| | | Post-Shock | Transient Peak | |
| | | | Stable Excitation | |
| | | | Stable Inhibition | |
| | | | None | |
| | Effect | Test | Statistic | |
| | Non-Classified (Emulated) Stable Excitation | Chi-Squared Goodness of Fit Test | $\chi^2(1) = 34.527$ | $P < 0.00001$ |
| | Non-Classified (Emulated) Transient Peak | Chi-Squared Goodness of Fit Test | $\chi^2(1) = 0.064$ | P=0.8 |
| | DLS (Emulated) Stable Excitation | Chi-Squared Goodness of Fit Test | $\chi^2(1) = 4.6545$ | P=0.03097 |
| | DLS (Emulated) Transient Peak | Chi-Squared Goodness of Fit Test | $\chi^2(1) = 32.41$ | $P < 0.00001$ |
| | DMS (Emulated) Stable Excitation | Chi-Squared Goodness of Fit Test | $\chi^2(1) = 0.16863$ | P=0.6813 |
| | DMS (Emulated) Transient Peak | Chi-Squared Goodness of Fit Test | $\chi^2(1) = 5.3774$ | P=0.0204 |
| 4C - Left | DLS/DMS (Total Ratio (Stable Excitation) | Permutation Test | | P=~0.79 |
| 4C - Right | DLS/DMS (Total Ratio (Transient Peak) | Permutation Test | | P=~0.013 |
| 4D | DLS/DMS Difference correlation with | Pearson Correlation | r = -0.597, t(9) = -2.23 | P=0.052 |

| Figure | Dependent Variable | Group | N | Mean | SD |
| --- | --- | --- | --- | --- | --- |
| 5C | Induced Grooming Length | Pre-Shock Excitation | 10 | 27.15 | 16.09 |
| | | Post-Shock Excitation | 10 | 20.55 | 16.20 |
| | | Pre-Shock Inhibition | 10 | 24.46 | 22.1 |
| | | Post-Shock Inhibition | 10 | 19.17 | 6.85 |
| | | Pre-Shock Drop-Only | 10 | 22.66 | 17.41 |
| | | Post-Shock Drop-Only | 10 | 31.27 | 13.2 |
| | | **Effect** | **Test** | **Statistic** | **p** |
| | | Trial_Type | Two-Way Within Anova | F(1,9)=0.042 | 0.853 |
| | | Measure | Two-Way Within Anova | F(2,18)=0.969 | 0.399 |
| | | Trial_type x Measure | Two-Way Within Anova | F(2,18)=4.67 | p = 0.0232 |
| | | Drop Only Vs Inhibition (Pre-Shock ) | Dunnet | t(18)=0.834 | P=0.622 |
| | | Drop Only Vs Excitation (Pre-Shock ) | Dunnet | t(18)=0.334 | P=0.922 |
| | | Drop Only Vs Inhibition (Post-Shock ) | Dunnet | t(18)=-2.855 | P=0.0196 |
| | | Drop Only Vs Excitation (Post-Shock ) | Dunnet | t(18)=-3.224 | P=0.0089 |
| 5D | ITI Grooming Length | Pre-Shock Excitation | 10 | 20.1 | 8.67 |
| | | Post-Shock Excitation | 10 | 14.7 | 3.16 |

| | | | | | |
|---|---|---|---|---|---|
| | | Pre-Shock Inhibition | 10 | 19.2 | 10.5 |
| | | Post-Shock Inhibition | 10 | 15.6 | 3.91 |
| | | Pre-Shock Drop-Only | 10 | 18.3 | 7.50 |
| | | Post-Shock Drop-Only | 10 | 21.6 | 6.42 |
| | | Effect | Test | Statistic | p |
| | | Trial_Type | Two-Way Within Anova | F(2,18)=2.034 | P=0.16 |
| | | Measure | Two-Way Within Anova | F(1,9)=0.919 | P=0.363 |
| | | Trial_type x Measure | Two-Way Within Anova | F(2,18)=3.537 | P=0.506 |
| | | Drop Only Vs Inhibition (Pre-Shock ) | Dunnet | t(18)=0.722 | P=0.697 |
| | | Drop Only Vs Excitation (Pre-Shock ) | Dunnet | t(18)=0.363 | P=0.909 |
| | | Drop Only Vs Inhibition (Post-Shock ) | Dunnet | T(18)=-3.452 | P=0.00539 |
| | | Drop Only Vs Excitation (Post-Shock ) | Dunnet | t(18)=-3.024 | P=0.0136 |
| 5E | Grooming Drop Initiation Delay | Pre-Shock Excitation | 10 | 7.72 | 4.01 |
| | | Post-Shock Excitation | 10 | 8.87 | 4.73 |
| | | Pre-Shock Inhibition | 10 | 9.73 | 5.57 |
| | | Post-Shock Inhibition | 10 | 10.2 | 6.81 |
| | | Pre-Shock Drop-Only | 10 | 7.71 | 2.97 |

| | | | |
|---|---|---|---|
| Post-Shock Drop-Only | 10 | 7.36 | 3.08 |

| Effect | Test | Statistic | p |
|---|---|---|---|
| Trial_Type | Two-Way Within Anova | F(2,18)=2.230.8 | P=0.136 |
| Measure | Two-Way Within Anova | F(1,9)=0.06 | P=0.812 |
| Trial_type x Measure | Two-Way Within Anova | F(2,18)=0.173 | P=0.843 |
| Drop Only Vs Inhibition (Pre-Shock ) | Dunnet | T(18)=0.005 | P=1 |
| Drop Only Vs Excitation (Pre-Shock ) | Dunnet | T(18)=0.983 | P=0.525 |
| Drop Only Vs Inhibition (Post-Shock ) | Dunnet | T(18)=1.126 | P=0.439 |
| Drop Only Vs Excitation (Post-Shock ) | Dunnet | T(18)=2.097 | P=0.0899 |

| Figure | Effect | Test | Statistics | p |
| --- | --- | --- | --- | --- |
| Supp1A | Weighted Correlation between Post-Stress Measures average induced grooming length and average Peak Height of Transient Peak Units responding to grooming termination | Weighted Pearson Correlation | r=0.525, t(14)=2.31 | P = 0.037 |
| Supp1B | Weighted Correlation between Post-Stress Measures average ITI length and average Peak Height of Transient Peak Units responding to grooming termination | Weighted Pearson Correlation | r=0.644, t(14)=3.15 | P=0.007 |
